# The J-Like Protein ARC6 Regulates Chloroplast FtsZ-Ring Assembly through Fine-tuning ARC3 Activity in Arabidopsis

**DOI:** 10.1101/2024.01.08.574726

**Authors:** Wenbin Du, Lingyan Cao, Yuelong Zhou, Shanelle Jackson, Maryam Naeem, Yue Yang, Jonathan M. Glynn, Katie J. Porter, Qian He, Jie Xu, Wanqi Liang, Katherine W. Osteryoung, Cheng Chen

## Abstract

Chloroplast division is initiated by the establishment of the stromal FtsZ ring (Z ring). Assembly and positioning of the Z ring are governed by the chloroplast Min system, which inhibits Z-ring formation everywhere but the middle of the chloroplast. ACCUMULATION AND REPLICATION OF CHLOROPLASTS3 (ARC3), the core component of this system, is a direct inhibitor of Z-ring assembly. Regulation of ARC3 activity is vital thus for chloroplast division. Here, we report that ARC6, which localizes on the chloroplast inner envelope membrane, interacts with ARC3 and acts upstream of ARC3 during chloroplast division. We show that the C-terminal MORN domain of ARC3, demonstrated previously to prevent ARC3-FtsZ interaction, binds to the J-like domain (JLD) of ARC6, enabling full-length ARC3 to interact with FtsZ proteins and activating the inhibitory activity of ARC3 on the assembly of FtsZ filaments. Overexpression of a JLD-deleted version of ARC6 causes disruption of Z-ring formation in an ARC3-dependent manner. Finally, we reveal that ARC6 recruits ARC3 to the middle of the chloroplast. Our findings suggest a model whereby ARC6 regulates the assembly and positioning of the Z ring through fine-tuning the inhibitory activity of ARC3 at the chloroplast division site.

**One Sentence Summary:** The chloroplast membrane protein ARC6 recruits ARC3 to the chloroplast division site and regulates the assembly of the FtsZ ring by fine-tuning ARC3 activity through its J-like domain.

The author responsible for distribution of materials integral to the findings presented in this article in accordance with the policy described in the Instructions for Authors (https://academic.oup.com/plcell/pages/General-Instructions) is: Cheng Chen.

## INTRODUCTION

Chloroplasts, the photosynthetic organelles, are derived from endosymbiosis when a cyanobacterium was engulfed by an ancient eukaryotic host cell about a billion years ago (Keeling, 2010; Zimorski et al., 2014; Martin et al., 2015). Thus, like the cyanobacterial endosymbiont, chloroplasts are surrounded by double membranes. Chloroplasts are responsible for the production of photosynthesis and for providing lipids, amino acids, and phytohormones (Cackett et al., 2022). During plant growth, the number of chloroplasts needs to increase to fulfill the ever-increasing demands for energy (Leech and Baker, 1983). Such an increase in chloroplast number is achieved through chloroplast division. Analogous to their prokaryotic ancestor, chloroplasts propagate from pre-existing organelles by binary fission (division in the middle) (Osteryoung and Pyke, 2014; Chen et al., 2018a).

Chloroplast division is carried out by the midplastid-localized macromolecular machinery (Osteryoung and Pyke, 2014; Yoshida, 2018). The major components of the division machinery are the contractile ring structures formed across the two chloroplast envelope membranes (Chen et al., 2018a). Among them, the stromal FtsZ ring (Z ring) is the first ring structure formed during chloroplast division (Miyagishima et al., 2001). The Z ring is assembled from the cytoskeletal GTPase protein FtsZ (TerBush et al., 2013; McQuillen and Xiao, 2020). In *Arabidopsis*, there are two FtsZ proteins, namely FtsZ1 and FtsZ2 (Osteryoung et al., 1998; Schmitz et al., 2009). Chloroplast FtsZ1 and FtsZ2 co-assemble into dynamic filaments and ring-like structures (Olson et al., 2010; Yoshida et al., 2016; Porter et al., 2021). The stromal Z ring is tethered to the inner envelope membrane both directly through the amphiphilic motif at the C-terminus of FtsZ1 (Liu et al., 2022) and indirectly through the interaction of FtsZ2 with the inner envelope membrane protein ARC6 (Maple et al., 2005; Johnson et al., 2013). The Z ring is highly dynamic, and it coordinates with the cytosolic DRP5B (also known as ARC5) ring to constrict (Gao et al., 2003; Miyagishima et al., 2003; Johnson et al., 2015; Yoshida et al., 2016). The inner and outer rings simultaneously constrict to squeeze the chloroplast membranes and eventually generate two daughter chloroplasts of approximately equal size.

The key step for chloroplast division is the positioning and subsequent constriction of the Z ring in the middle of the chloroplast. This is governed by the chloroplast Min system, which is a negative regulatory mechanism inhibiting the assembly of the Z ring at non-division sites in order to allow Z-ring formation only in the middle of the chloroplast (Osteryoung and Pyke, 2014; Chen et al., 2018a). The chloroplast Min system consists of ARC3, MinD1, MinE1, and MCD1 (Colletti et al., 2000; Maple et al., 2002; Maple et al., 2007; Nakanishi et al., 2009). Among them, ARC3 is the central player, acting as the direct inhibitor of Z-ring assembly (TerBush and Osteryoung, 2012; Zhang et al., 2013). There are multiple Z rings in the enlarged chloroplasts of *arc3* mutants, while Z-ring formation is abolished in transgenic plants overexpressing ARC3 (Glynn et al., 2007; Zhang et al., 2013).

The inhibitory activity of ARC3 on FtsZ assembly is regulated by its C-terminal MORN domain (Figure 1A) (Zhang et al., 2013). Previous studies demonstrated that the MORN domain prevents ARC3 from interacting with the FtsZ proteins (Zhang et al., 2013), implying the existence of auto-inhibition in terms of the activity of ARC3 on the assembly of the FtsZ filaments. Previous studies reported that PARC6, an inner envelope membrane protein, binds ARC3 through the MORN domain, and such binding activates the inhibitory activity of ARC3 on the assembly of the FtsZ filaments (Glynn et al., 2009; Zhang et al., 2016; Chen et al., 2019). ARC3 localizes both diffusely in the stroma and as a ring-like structure at the division site (Chen et al., 2019). It was proposed that the diffuse ARC3 is mainly responsible for inhibiting Z-ring formation at non-division sites. Regarding the midplastid ARC3, it has been suggested that it will enhance the dynamics of the Z ring and facilitate the constriction of chloroplasts (Johnson et al., 2015; Chen et al., 2019). Despite these findings, it still remains elusive how the activity of ARC3 is regulated, specifically at the division site, since the activity of ARC3 there must be precisely controlled in order to simultaneously allow the establishment of the Z ring and to promote the dynamics of the Z ring during the constriction and division of chloroplast.

**Figure 1.**
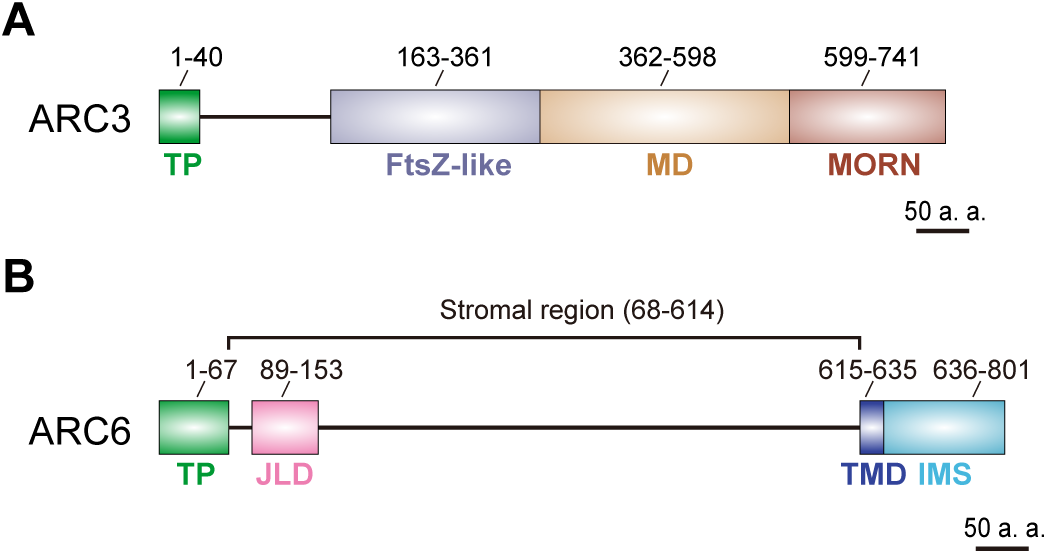
Schematic depiction of ARC3 and ARC6 domain structures. Schematic structural features of **(A)** ARC3 and **(B)** ARC6 from *Arabidopsis thaliana*. TP, predicted chloroplast transit peptide; FtsZ-like, FtsZ-like domain; MD, middle domain; MORN, Membrane Occupation and Recognition Nexus domain; JLD, J-like domain; TMD, transmembrane domain; IMS, intermembrane space region; a. a., amino acid.

ARC6 has been shown to play an indispensable role in chloroplast division, as evidenced by the severe chloroplast division defect in the *arc6* mutant plant. The mesophyll cells of *arc6* typically harbor only 1∼2 giant chloroplasts per cell (Pyke et al., 1994). In addition, the formation of the Z ring is totally disrupted, and only dots or patches of the FtsZ filaments are observed in the *arc6* mutant (Vitha et al., 2003), indicating that ARC6 is a positive regulator for Z-ring formation. This could be due to the prevention of the disassembly of GDP-bound FtsZ2 protein by ARC6, as suggested by a recent study (Sung et al., 2018). Given that both FtsZ proteins are still able to assemble into filaments even in the absence of ARC6 *in vitro* (Olson et al., 2010; Porter et al., 2021) or *ex vivo* (TerBush and Osteryoung, 2012; TerBush et al., 2016; Yoshida et al., 2016; TerBush et al., 2018), it could not be totally accountable for the disruption of Z-ring formation in the *arc6* mutant. ARC6 has been shown to bridge MCD1 to interact with FtsZ2, which may allow MCD1 to recognize the membrane-tethered Z ring and ultimately to regulate the positioning of the division machinery (Chen et al., 2018b). A recent study reported that ARC6 directly interacts with MinD1 (Zhang et al., 2021).The biological significance of such an interaction, however, is unsolved. Nevertheless, the molecular mechanisms by which ARC6 is used to regulate chloroplast division, particularly Z-ring assembly and positioning, are still lacking.

ARC6 is a descendant of the cyanobacterial protein Ftn2 (also known as ZipN) (Vitha et al., 2003). In addition to the transmembrane domain, there is a J-like domain downstream of the N-terminal predicted chloroplast transit peptide of ARC6 (Figure 1B). The J domain was originally designated for the DNAJ proteins, which serve as cochaperones of Hsp70 proteins (Pulido and Leister, 2018; Tamadaddi et al., 2022).

The J domain is responsible for DNAJ proteins to interact with Hsp70s (Pulido and Leister, 2018), which results in the stimulation of Hsp70 ATPase activity and may provide substrate specificity to Hsp70-mediated processes. However, the J-like domain of ARC6 is not a canonical J domain since it lacks the conserved tripeptide HPQ found in DNAJ proteins (Pulido and Leister, 2018). The J-like domain of ARC6 is not required for interaction with FtsZ2 (Maple et al., 2005; Glynn et al., 2009; Zhang et al., 2016). In contrast, it is required for interaction with CJD, a J-like protein involved in the regulation of the fatty acid composition of chloroplast lipids, though the biological significance is unknown (Ajjawi et al., 2011). Thus, the function of the J-like domain of ARC6, particularly in chloroplast division, remains elusive.

Here, we investigated the functional relationship between ARC6 and ARC3 and characterized the underlying molecular mechanisms by which ARC6 regulates ARC3 during chloroplast division. Our results reveal that ARC6 directly binds to ARC3 and enables full-length ARC3 to interact with FtsZ proteins. We further demonstrated that ARC6 activates the inhibitory activity of ARC3 on the assembly of FtsZ filaments in a heterologous system. The activation of ARC3 by ARC6 is fine-tuned by the J-like domain of ARC6, allowing the establishment of the Z ring at the division site. Finally, we show that the midplastid-localized ARC3 is recruited by ARC6. Our findings unveil novel functions of ARC6 during chloroplast division and advance the understanding of the regulatory mechanisms of the formation and positioning of the Z ring at the division site.

## RESULTS

### ARC6 Is Required for ARC3 to Function in Chloroplast Division

It has been reported that ARC6 and ARC3 are purified together in a complex containing FtsZ proteins in Arabidopsis (McAndrew et al., 2008), suggesting a possible functional dependency. To investigate the functional relationship between *ARC6* and *ARC3* during chloroplast division, we generated a double mutant by crossing the *arc6-5* and *arc3-2* single mutants. Unlike wild-type Col-0, which harbored plenty of relatively small chloroplasts within the mesophyll cells (Figure 2, A and E), both *arc3-2* and *arc6-5* single mutants contained a reduced number while enlarged chloroplasts (Figure 2, B, C and E), indicating a defect in chloroplast division. In line with prior reports, the chloroplast division defect was more profound in *arc6-5*, which harbored only 1∼2 giant chloroplasts within mesophyll cells, as compared to the *arc3-2* mutant (Figure 2, B, C and E) (Maple et al., 2007; Ajjawi et al., 2011; Zhang et al., 2013). We found that both the morphology of chloroplasts and the plant growth phenotype observed in the *arc3-2 arc6-5* mutant were similar to those detected in *arc6-5* (Figure 2, B to E; Supplemental Figure 1).

**Figure 2.**
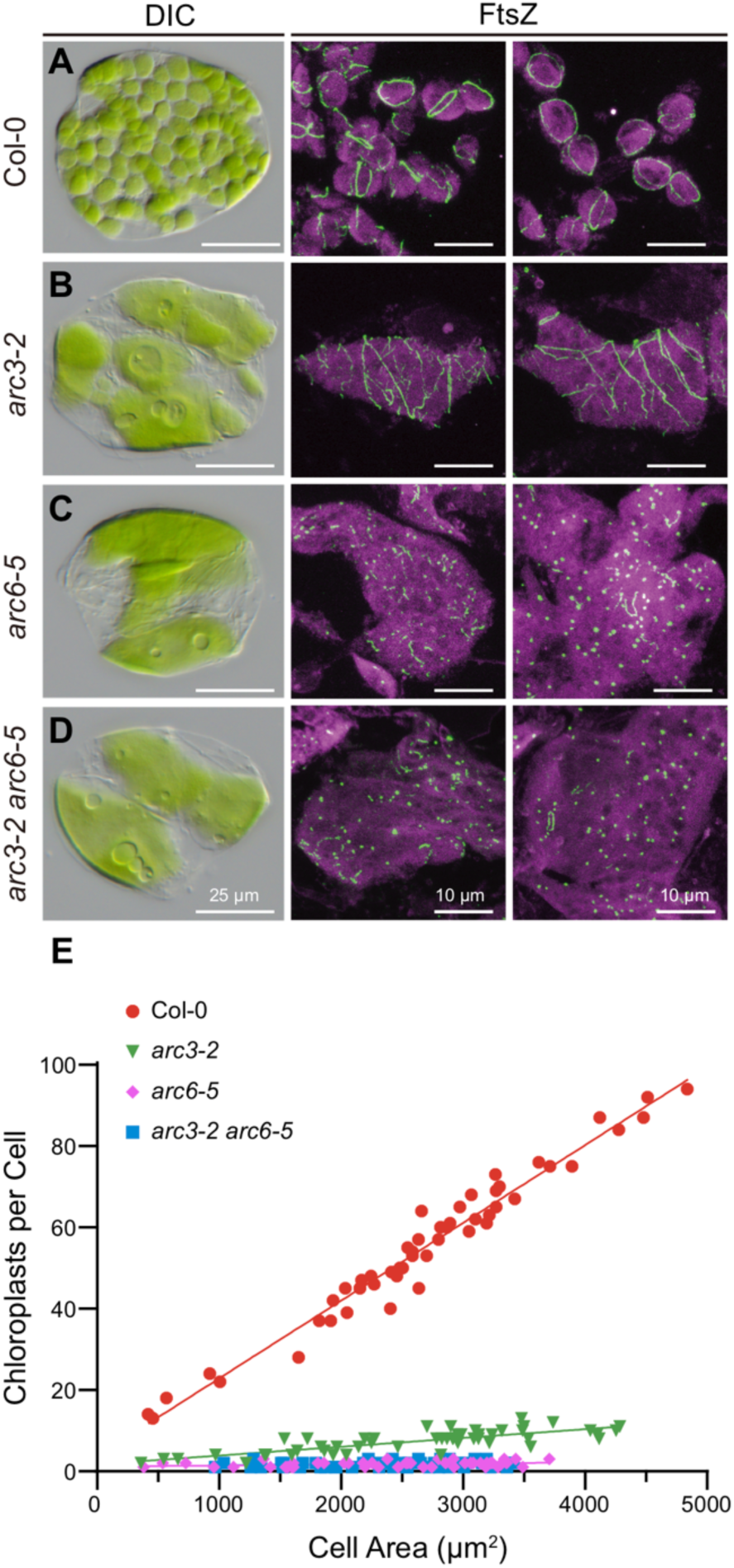
ARC6 is required for ARC3 function in chloroplast division. **(A-D)** Chloroplast morphology (left panels) and FtsZ localization (middle and right panels) in mesophyll cells of **(A)** wild-type Col-0, **(B)** *arc3-2*, **(C)** *arc6-5*, and **(D)** *arc3-2 arc6-5* mutant plants. Chloroplast morphology and FtsZ localization were observed using differential interference contrast (DIC) microscopy and immunofluorescence staining of FtsZ2-1, respectively. The middle and right panels are merged images of immunostained FtsZ2-1 (green) and autofluorescence of chlorophyll (magenta). Scale bars are 25 μm (left panels) and 10 μm (middle and right panels), respectively. **(E)** Quantitative analysis of chloroplast number versus mesophyll cell size (*n* = 50) in the designated genotypes from **(A-D)** The computed slopes of the best-fit lines are: Col-0, 0.0191 (R^2^ = 0.965); *arc3-2*, 0.0022 (R^2^ = 0.635); *arc6-5*, 0.0002 (R^2^ = 0.102); *arc3-2 arc6-5*, 0.0002 (R^2^ = 0.024).

To determine the assembly and positioning of the Z ring in these mutant plants, we further conducted immunofluorescence staining. We found that a single Z ring was formed in the middle of chloroplasts from the wild-type Col-0 (Figures 2A, middle and right panels). In contrast, multiple Z rings across the whole chloroplast were detected in the *arc3-2* mutant (Figure 2B, middle and right panels) (Zhang et al., 2013). As previously reported for an allelic mutant of *ARC6* (*arc6-1*) (Vitha et al., 2003), only short FtsZ fragments or puncta were observed in the *arc6-5* mutant (Figures 2C, middle and right panels). Similar to *arc6-5*, we observed only short FtsZ fragments, but not the intact Z ring, in the *arc3-2 arc6-5* double mutant (Figure 2D, middle and right panels). These results suggest that ARC6 acts upstream of ARC3 during chloroplast division.

### ARC6 Interacts with ARC3

Given the functional relationship between ARC6 and ARC3, we were prompted to ask whether they interact directly. To this end, we adopted the yeast two-hybrid (Y2H) assay. Since the N-terminus of ARC6 faces the stroma where ARC3 is localized (Vitha et al., 2003), we generated a truncated version of ARC6 lacking the predicted transit peptide, ARC6_68-614_ (Figure 1B), and tested its interaction with various versions of ARC3. As revealed in Figure 3A, ARC6_68-614_ interacted with full-length ARC3 (ARC3_41-741_) (Figure 3A, row 1) but not with a version of ARC3 missing the MORN domain (ARC3_41-598_) (Figure 3A, row 2). Interestingly, neither ARC3_Δ630-675_ nor ARC3_6G to A_ interacted with ARC6_68-614_ (Figure 3A, rows 4 and 5), demonstrating that the C-terminal MORN domain of ARC3 is critical for its interaction with ARC6 since the function of the MORN domain was disrupted within both ARC3 derivatives (Chen et al., 2019). Indeed, we showed that ARC6 directly binds to the MORN domain of ARC3 (ARC3_598-741_) (Figure 3A, row 3). In line with prior reports, ARC6 interacted with FtsZ2 but not with FtsZ1, which served as a positive and negative control, respectively (Figure 3A, rows 7 and 6) (Maple et al., 2005; Glynn et al., 2009; Schmitz et al., 2009).

**Figure 3.**
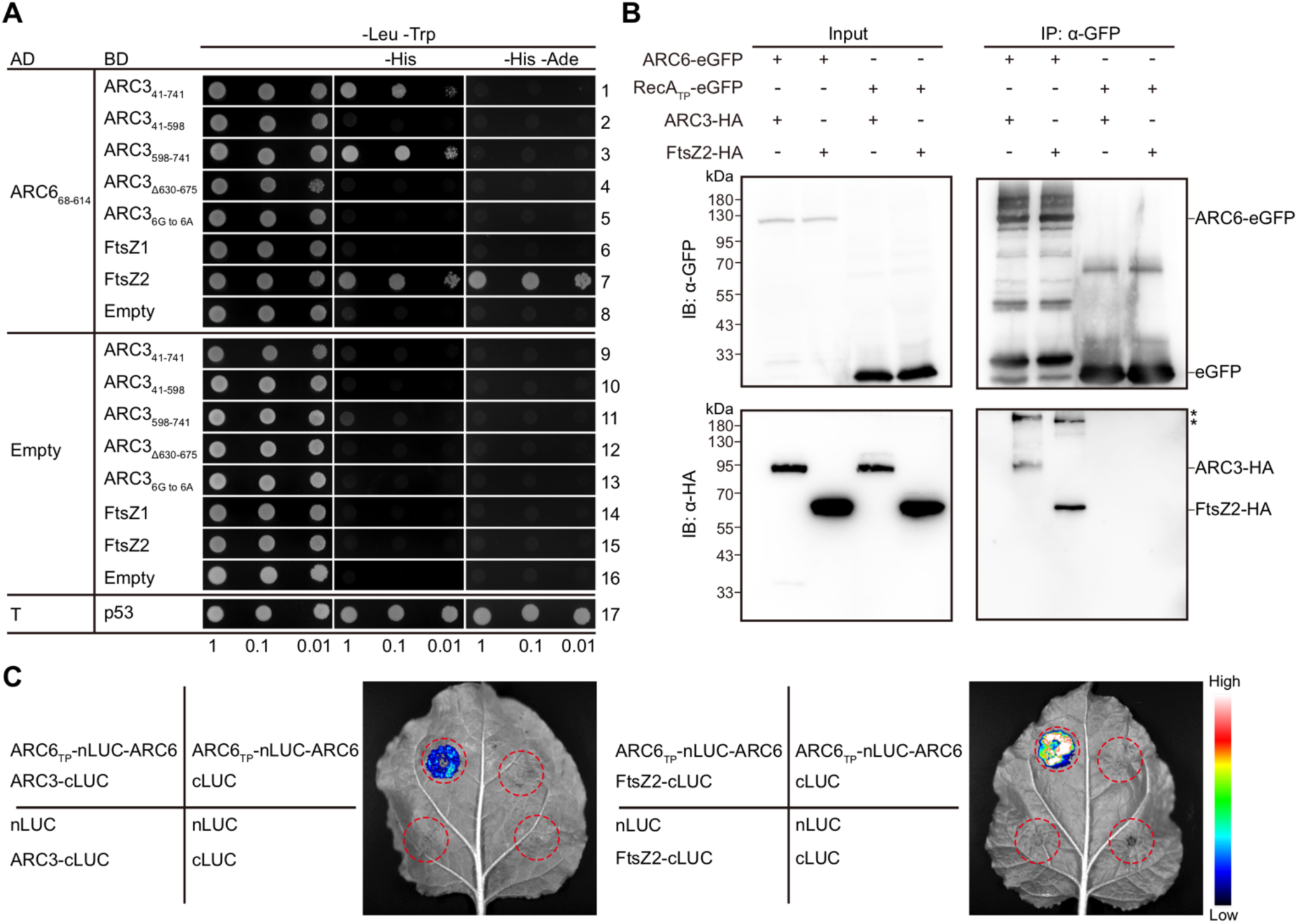
ARC6 interacts with ARC3. **(A)** Yeast two-hybrid (Y2H) assays of ARC6_68-614_ with ARC3 derivatives and FtsZ proteins. Constructs were expressed from the pGADT7 (AD) and pGBKT7 (BD) vectors in Y2HGold cells. Transformed cells were grown on a selective medium lacking leucine and tryptophan (–Leu –Trp). Interactions were determined based on the activation of the *HIS3* and *ADE2* reporter genes, as indicated by the growth on medium lacking histidine (–His) alone or both histidine and adenine (–His –Ade). The interaction between Simian Virus-40 large-T antigen (T) and p53 was used as a positive control (row 17). Empty vectors were used as negative controls. Transformed cells were grown on –Leu –Trp and –Leu –Trp –His medium for three days, and five days on –Leu –Trp –His –Ade medium. Dilutions from the same initial culture are denoted at the bottom. Three independent replicates yielded similar results. **(B)** Co-immunoprecipitation (Co-IP) assays of ARC6-eGFP with ARC3-6×HA and FtsZ2-6×HA in *Nicotiana benthamiana*. The transit peptide (TP) of RecA, a stroma-localized protein (Kohler et al., 1997), was fused to the N-terminus of eGFP to generate RecA_TP_-eGFP fusion protein (Chen et al., 2018b), which served as control. Total proteins were extracted 48 h after agroinfiltration, and samples were separated on 10% SDS-PAGE. Asterisks indicate potential protein complexes formed by ARC6-ARC3 and ARC6-FtsZ2, respectively. **(C)** Split-luciferase complementation (split-LUC) assays of ARC6 with ARC3 and FtsZ2. Samples were observed 48 h after agroinfiltration into the *N. benthamiana* leaves. N-terminal luciferase (nLUC) and C-terminal luciferase (cLUC) were used as negative controls. Red dashed circles indicated the region of infiltration. Assays in **(B)** and **(C)** were replicated three times with similar results.

To explore whether ARC6 interacts with ARC3 *in planta*, we performed the Co-immunoprecipitation (Co-IP) assay using tobacco plants. The results showed that ARC3-6×HA could be co-precipitated with the ARC6-eGFP fusion protein (Figure 3B). As a positive control, we showed that FtsZ2-6×HA interacted with ARC6-eGFP in this assay (Figure 3B). To further validate the ARC6-ARC3 interaction *in vivo*, we implemented a split-luciferase complementation (split-LUC) assay in tobacco. Consistent with the results obtained from the Y2H and Co-IP assays, we detected a direct interaction between ARC6 and ARC3 in the split-LUC assay (Figure 3C). As a positive control, the ARC6-FtsZ2 interaction was observed as well (Figure 3C). Thus, our data demonstrate that ARC6 directly binds to ARC3, presumably through the C-terminal MORN domain of ARC3.

### ARC6-ARC3 Interaction Is Regulated by the J-like Domain of ARC6

ARC6 contains a conserved J-like domain, but its biological function remains elusive. Phylogenetic analysis revealed that ARC6 proteins from various species harbor a conserved PPQ motif within the J-like domain (Supplemental Figure 2). Thus, the ARC6 J-like domain is not a *bona fide* J domain since it lacks the conserved HPQ motif of the conventional J domain (Vitha et al., 2003). Accordingly, we found that the J-like domain of ARC6 did not interact with chloroplastic Hsp70 proteins (Supplemental Figure 3). To explore the function of the ARC6 J-like domain, we constructed ARC6 variants lacking either the intact J-like domain (ARC6_ΔJLD_) or only the PPQ motif (ARC6_ΔPPQ_), and tested their interaction with various ARC3 derivatives. Our Y2H results showed that the ARC6_ΔJLD_ not only interacted with full-length ARC3 and the MORN domain of ARC3 (Figure 4A, rows 7 and 9), but also with the MORN-truncated version of ARC3 (Figure 4A, row 8). Moreover, ARC6_ΔJLD_ interacted more strongly with full-length ARC3 and the MORN domain of ARC3 compared to ARC6 (Figure 3A, rows 1 and 3, right panels), as evidenced by the growth of the transformed yeast cells on the more stringent –His –Ade selective plates (Figure 4A, rows 7 and 9, right panels). We further found that the J-like domain of ARC6 could directly bind to the ARC3 MORN domain (Figure 4A, row 21).

**Figure 4.**
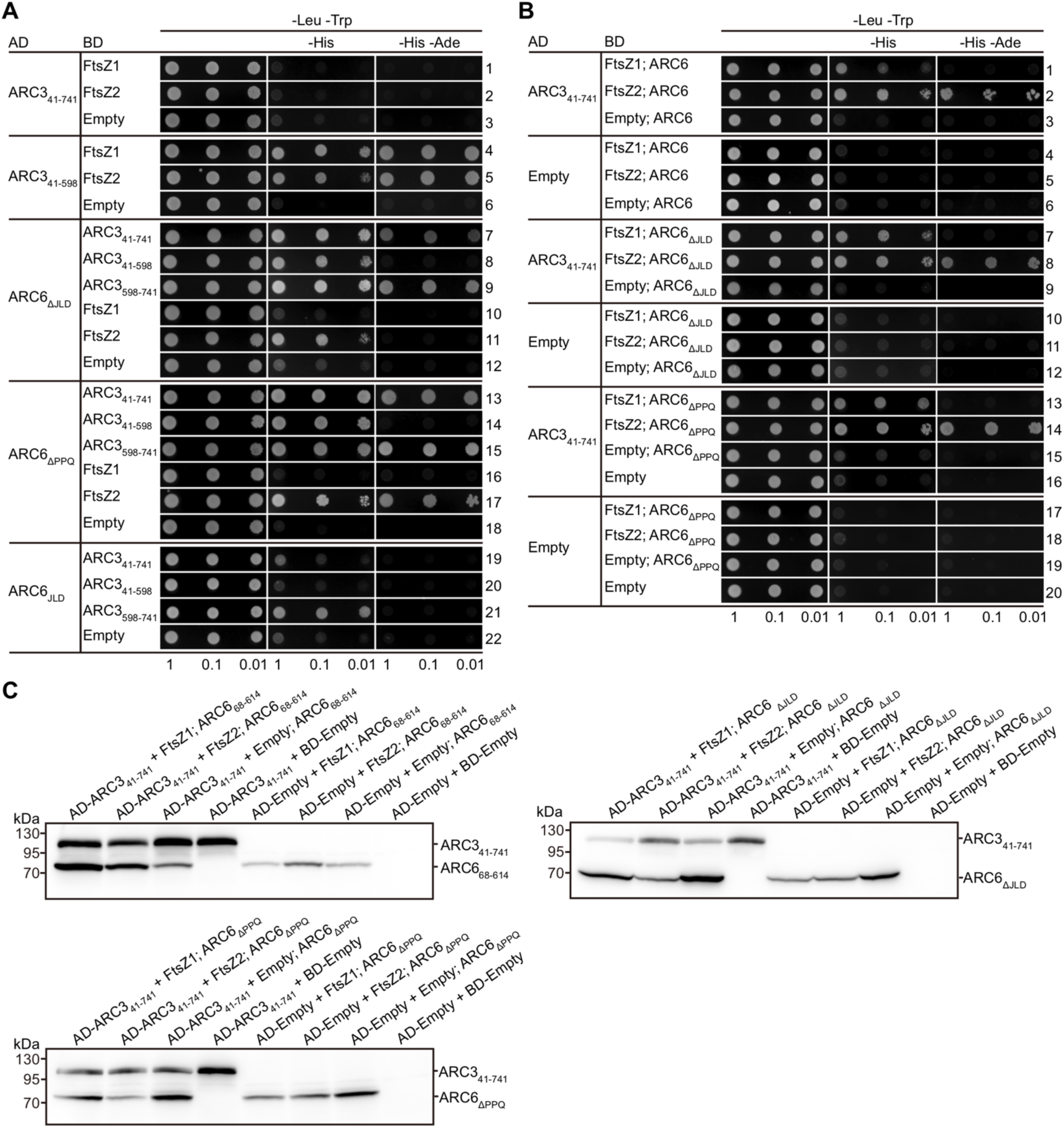
ARC6 enables full-length ARC3 to interact with FtsZ proteins. **(A)** Y2H assays to investigate interactions between ARC3 and FtsZ proteins and between ARC6 derivatives and ARC3. ARC6_ΔJLD_, ARC6_68-614_ without the JLD domain; ARC6_ΔPPQ_, ARC6_68-614_ without the tripeptide PPQ; ARC6_JLD_, ARC6_89-153_. **(B)** Y3H assays to test for interactions between full-length ARC3 (ARC3_41-741_) and FtsZ proteins in the presence of ARC6 derivatives. Vectors and assays are as described in Figure 3A except that in vectors expressing ARC6 derivatives, ARC6 derivatives were not fused to the GAL4 activation domain or binding domain. The ARC6 expression cassettes were inserted into the pGBKT7 vectors (BD) expressing the BD-FtsZ1, BD-FtsZ2, and BD-Empty constructs (see Supplemental Figure 4). Transformed cells in **(A)** and **(B)** were grown on –Leu –Trp and –Leu –Trp –His medium for three days, and five days on –Leu –Trp –His –Ade medium. Dilutions from the same initial culture are indicated at the bottom. Assays in **(A)** and **(B)** were replicated three times with similar results. **(C)** Immunoblot assays of transformed yeast cells expressing the indicated constructs. Total proteins were separated on 10% SDS-PAGE gels, and membranes were probed with an anti-HA antibody, given that all ARC6 derivatives and ARC3 fusion proteins were HA-tagged. The predicted protein molecular weights were: AD-ARC3_41-741_, 96 kDa; ARC6_68-614_, 66 KDa; ARC6_ΔJLD_, 59 kDa; ARC6_ΔPPQ_, 66 kDa. AD-ARC3_41-741_ ran slightly larger than predicted.

Analogous to ARC6_ΔJLD_, ARC6_ΔPPQ_ exhibited a similar interaction pattern with the various ARC3 derivatives (Figure 4A, rows 13 to 18). Intriguingly, the interaction between ARC6 and FtsZ2 was reduced, but not abolished, by the truncation of the J-like domain (Figure 4A, row 11), whereas it was not affected if only the PPQ motif was deleted (Figure 4A, row 17), suggesting that the entire J-like domain contributes to the ARC6-FtsZ2 interaction. Taken together, these data suggest that the unconventional J-like domain of ARC6 regulates both ARC6-ARC3 and ARC6-FtsZ2 interactions.

### ARC6 Enables Full-length ARC3 to Interact with FtsZ Proteins

The MORN domain has previously been shown to prevent ARC3 from interacting with FtsZ proteins. In line with the prior reports, ARC3_41-741_, the full-length ARC3, did not interact with either FtsZ1 or FtsZ2 due to the presence of the MORN domain (Figure 4A, rows 1 and 2) (Maple et al., 2007; Zhang et al., 2013). In contrast, ARC3_41-598_, the MORN-truncated version of ARC3, interacted with both FtsZ proteins (Figure 4A, rows 4 and 5), and the transformed yeast cells even grew on the more stringent –His –Ade selective plates as shown in the Y2H assay (Figure 4A, rows 4 to 6, right panels).

Based on our finding that ARC6 directly binds to the MORN domain of ARC3, we hypothesized that the binding of ARC6 to ARC3 may sequester the MORN domain, thereby allowing ARC3 to interact with FtsZ proteins. To test this hypothesis, we conducted the yeast three-hybrid (Y3H) assay (Chen et al., 2019). In contrast to the results obtained from Y2H, our Y3H data showed that ARC3_41-741_ interacted with both FtsZ proteins when ARC6 was included (Figure 4B, rows 1 and 2). Intriguingly, ARC3_41-741_ bound more strongly with FtsZ2 than with FtsZ1 in the presence of ARC6, as evidenced by the growth of the transformed yeast cells on –His –Ade plates for FtsZ2 (Figure 4B, rows 1 and 2, right panels). This could be due, at least in part, to the ARC6-FtsZ2 interaction (Figure 3A, row 7), whereas ARC6 does not interact with FtsZ1 (Figure 3A, row 6) (Maple et al., 2005). The obtained positive interactions resulted only from interaction between ARC3_41-741_ and FtsZ proteins since the ARC6 expression cassette in the Y3H assay contained neither the GAL4 activation domain (AD) nor the binding domain (BD) and thus was unable to activate expression of the reporter genes (Supplemental Figure 4). This was verified when vectors encoding AD-Empty and BD-FtsZ2; ARC6 were cotransformed, and no growth of the transformed yeast cells on the selective plates could be observed in Y3H (Figure 4B, row 5), despite the fact that AD-ARC6 interacts with BD-FtsZ2 in standard Y2H assays (Figure 3A, row 7). The expression of ARC6 and ARC3 in these Y3H assays was validated by Western blot (Figure 4C). Altogether, these results demonstrate that ARC6 enables full-length ARC3 to interact with FtsZ proteins.

We further adopted the Y3H assay to determine the effect of the J-like domain on the ARC6-ARC3-FtsZ interaction. Our data revealed that both ARC6_ΔJLD_ and ARC6_ΔPPQ_ enabled full-length ARC3 to interact with FtsZ proteins (Figure 4B, rows 7, 8, 13 and 14). Moreover, both derivates were more efficient in promoting the ARC3-FtsZ interactions when compared to wild-type ARC6 (Figure 4B, rows 7, 8, 13 and 14; Supplemental Figure 5). Intriguingly, ARC6_ΔPPQ_ exhibited a stronger ability to enable the ARC3-FtsZ interactions than did ARC6_ΔJLD_ (Supplemental Figure 5, rows 7, 8, 13 and 14). Western blot analysis validated the expression of ARC6_ΔJLD_ and ARC6_ΔPPQ_ as well as ARC3 fusion proteins in these Y3H assays (Figure 4C). These data suggest that the J-like domain of ARC6 ultimately regulates the ARC3-FtsZ interactions enabled by ARC6.

### ARC6 Activates the Inhibitory Activity of Full-length ARC3 on the Assembly of FtsZ Filaments

Given that ARC6 enables full-length ARC3 to directly interact with FtsZ proteins, we next investigated whether ARC6 could further activate the inhibitory activity of ARC3 on the assembly of FtsZ filaments. Towards addressing this, we adopted the heterologous yeast *Schizosaccharomyces pombe* to investigate the assembly of chloroplast FtsZ filaments (TerBush and Osteryoung, 2012; TerBush et al., 2016; TerBush et al., 2018). *S. pombe* lacks endogenous FtsZ and its regulators, which makes it a valuable system to address our hypothesis. Previous studies reported that ARC6 colocalizes with FtsZ2 filaments when coexpressed in *S. pombe* yeast cells (TerBush et al., 2016; Sung et al., 2018). To exclude the potential effect of ARC6 on FtsZ2 due to their direct interaction, we used FtsZ1 instead in this assay. We generated constructs encoding ARC3-mCerulean, ARC6-mRuby2, and FtsZ1-mVenus fusion proteins and expressed them in *S. pombe* cells. As previously shown (TerBush et al., 2016; Chen et al., 2019), FtsZ1-mVenus assembled into filaments (Figure 5A), while ARC6-mRuby2 was diffuse when expressed alone in *S. pombe* cells (Figure 5B). Consistent with prior reports (TerBush and Osteryoung, 2012; Zhang et al., 2013; Chen et al., 2019), we observed both aggregates and diffuse signals formed by ARC3_41-741_-mCerulean (Figure 5C), but only diffuse signals for ARC3_41-598_-mCerulean (Figure 5D). In line with the ARC3-FtsZ interaction data obtained from the above Y2H assays, ARC3_41-741_-mCerulean did not (Figure 5E), whereas ARC3_41-598_-mCerulean did (Figure 5F), inhibit the assembly of FtsZ1 filaments, as evidenced by the decrease in filament length and increase in filament number of the assembled FtsZ1 filaments. As expected, ARC6 alone did not interfere with the assembly of FtsZ1 filaments due to a lack of interaction between them (Figure 5G).

**Figure 5.**
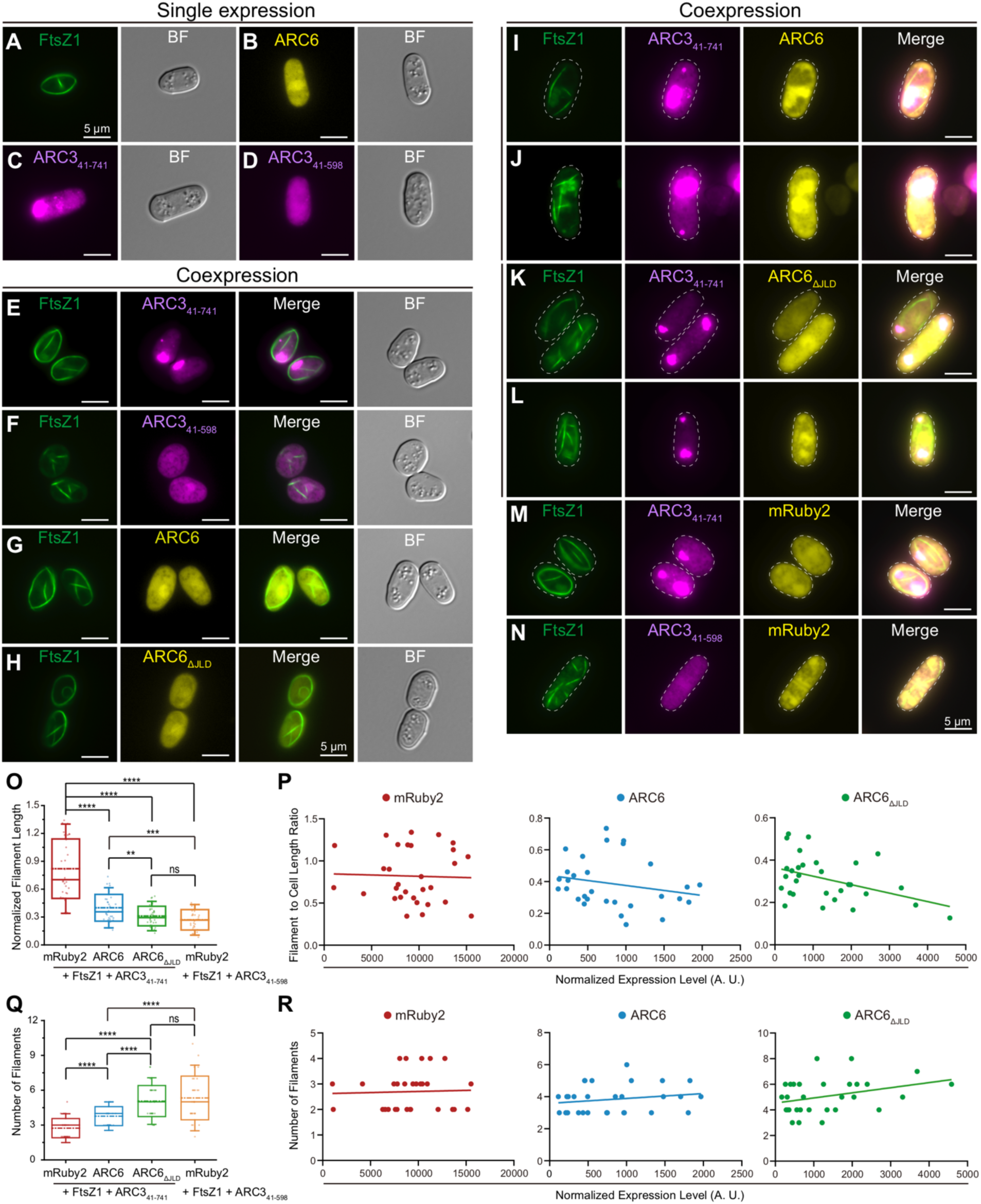
Activation of full-length ARC3 by ARC6 inhibits assembly of FtsZ1 filaments in *S. pombe*. The epifluorescence and DIC micrographs of transformed *S. pombe* cells expressing the indicated proteins are shown. In epifluorescence images, mVenus, mCerulean, and mRuby2 signals are falsely colored green, magenta, and yellow, respectively. BF, bright field. Bars = 5 μm. **(A-D)** Single expression of **(A)** FtsZ1-mVenus, **(B)** ARC6_68-614_-mRuby2, **(C)** ARC3_41-741_-mCerulean, and **(D)** ARC3_41-598_-mCerulean. **(E-H)** Coexpression of FtsZ1-mVenus with **(E)** ARC3_41-741_-mCerulean, **(F)** ARC3_41-598_-mCerulean, **(G)** ARC6_68-614_-mRuby2, **(H)** ARC6_ΔJLD_-mRuby2, **(I, J)** ARC3_41-741_-mCerulean + ARC6_68-614_-mRuby2, **(K, L)** ARC3_41-741_-mCerulean + ARC6_ΔJLD_-mRuby2, **(M)** ARC3_41-741_-mCerulean + mRuby2, and **(N)** ARC3_41-598_-mCerulean + mRuby2. Outlines of the imaged cells from **(I-N)** are indicated by white dashed lines. **(O-R)** Quantitative analysis of ARC3 derivatives on the assembly of FtsZ1 filaments in the presence of ARC6_68-614_-mRuby2 (*n* = 30 cells), ARC6_ΔJLD_-mRuby2 (*n* = 30 cells), or mRuby2 (*n* = 30 cells). **(O)** The length and **(Q)** number of FtsZ1 filaments were calculated to determine the inhibitory effect imposed by ARC3. In **(O)**, filament length was normalized to cell length, and the average of the two shortest filaments within the cell was presented. Error bars are SD. \*\*\*\**P* < 0.0001; \*\*\**P* < 0.001; \*\**P* < 0.01 as determined by the *t* test. ns, not significant. **(P)** The length of FtsZ1 filaments in **(O)** was plotted against the protein expression level of mRuby2 (left, red), ARC6_68-614_-mRuby2 (middle, blue), or ARC6_ΔJLD_-mRuby2 (right, green). The slopes of the best-fit lines are: mRuby2, -3e-6 (R^2^ = 0.001); ARC6_68-614_-mRuby2, -6e-5 (R^2^ = 0.048); ARC6_ΔJLD_-mRuby2, -4e-5 (R^2^ = 0.184). **(R)** The number of FtsZ1 filaments in **(Q)** was plotted against the protein expression level of mRuby2 (left, red), ARC6_68-614_-mRuby2 (middle, blue), or ARC6_ΔJLD_-mRuby2 (right, green). The slopes of the best-fit lines are: mRuby2, 9e-6 (R^2^ = 0.002); ARC6_68-614_-mRuby2, 3e-4 (R^2^ = 0.037); ARC6_ΔJLD_-mRuby2, 4e-4 (R^2^ = 0.114). The protein expression level in **(P, R)** was indicated by the total fluorescence intensity normalized to the cell area. A. U., arbitrary unit.

To determine the effect of ARC6 on the inhibitory activity of ARC3 on the assembly of FtsZ filaments, we cotransformed *FtsZ1-mVenus* with constructs expressing *ARC6-mRuby2* and *ARC3-mCerulean*. Our data showed that the assembly of FtsZ1 filaments was not affected when ARC3_41-741_-mCerulean was coexpressed with the fluorescent tag protein mRuby2 (Figure 5M), which served as a control. However, we found that the assembly of FtsZ1 filaments was significantly inhibited by ARC3_41-741_-mCerulean when ARC6-mRuby2 was introduced in the yeast cells (Figure 5, I and J; Supplemental Figure 6), as demonstrated by the decrease in the length of the filaments and increase in the number of filaments within the yeast cells (Figure 5, O and Q). The inhibition was positively correlated with the protein level of ARC6-mRuby2 (Figure 5, P and R, middle panels), but not with the level of mRuby2 (Figure 5, P and R, left panels), demonstrating that full-length ARC3-mediated inhibition of FtsZ1 assembly was genuinely imposed by ARC6. As expected, ARC3_41-598_-mCerulean inhibited the assembly of FtsZ1 filaments when coexpressed with mRuby2, which served as an additional control (Figure 5, N, O and Q). Intriguingly, we noticed that the MORN-truncated version of ARC3 was more efficient in inhibiting the assembly of FtsZ1 filaments than the full-length ARC3-ARC6 complex (Figure 5, I, J, N, O and Q). Taken together, these data provide direct evidence that ARC6 activates the inhibitory activity of ARC3 on the assembly of FtsZ filaments. Moreover, our data suggest that ARC6 not only activates ARC3, but also constrains it enough to allow the assembly of FtsZ filaments to occur to some extent.

### ARC6 Acts through Its J-like Domain to Fine-tune the Inhibitory Activity of ARC3 on the Assembly of FtsZ Filaments

The observation that the J-like domain of ARC6 regulates the ARC6-ARC3 interaction prompted us to investigate whether ARC6 could fine-tune the inhibitory activity of ARC3 on the assembly of FtsZ filaments through this domain. To this end, we first examined the assembly of FtsZ1 filaments in the presence of full-length ARC3 and ARC6_ΔJLD_ using the above heterologous yeast system. Our results showed that ARC6_ΔJLD_-mRuby2 increased the inhibitory activity of ARC3_41-741_-mCerulean on FtsZ1-mVenus filament assembly in *S. pombe* cells (Figure 5, K, L, O and Q), as indicated by the more significant changes in filament length and number in the transformed yeast cells (Figure 5, O and Q). Such inhibition was positively correlated with the protein level of ARC6_ΔJLD_-mRuby2 (Figure 5, P and R, right panels).

These findings led us to further investigate the function of the J-like domain of ARC6 *in vivo*. To this end, we generated constructs expressing *ARC6-mNeonGreen-3×HA* (*ARC6-mNG-3×HA*) and *ARC6_ΔJLD_-mNG-3×HA*, both driven by the *35S* promoter, and expressed them in Arabidopsis. Consistent with a prior report (Vitha et al., 2003), we found that overexpression of ARC6-mNG-3×HA in wild-type Col-0 caused multiple Z-ring formation and thus led to chloroplast division defects in transgenic plants (Supplemental Figure 7). In contrast, we observed that overexpression of ARC6_ΔJLD_-mNG-3*×*HA inhibited the assembly and formation of the Z ring, which resulted in the defect of chloroplast division in transgenic plants (Figure 6, A and B; Supplemental Figure 8). Interestingly, we observed the formation of mini-Z rings in transgenic plants overexpressing ARC6_ΔJLD_-mNG-3*×*HA (Figure 6A), resembling those observed in transgenic plants overexpressing ARC3 (Zhang et al., 2013; Chen et al., 2019). To determine whether ARC6_ΔJLD_-caused inhibition of FtsZ assembly depends on ARC3, we transformed *35S_pro_:ARC6_ΔJLD_-mNG-3*×*HA* into the *arc3-2* mutant. We found that the chloroplast division defect in these transgenic plants (Figure 6, C and D, left panels) resembled that in the non-transformed *arc3-2* mutant plants (Figure 2B, left panel). Immunofluorescence staining results showed that there were multiple Z rings in *arc3-2* transformed with *35S_pro_:ARC6_ΔJLD_-mNG-3*×*HA* (Figure 6, C and D, right panels), suggesting that ARC6_ΔJLD_ acted through ARC3 to impose the inhibition on Z-ring assembly and formation. Expression of the ARC6_ΔJLD_ fusion protein was at a comparable level in wild-type Col-0 and *arc3-2* mutant transgenic plants, as detected by Western blot (Figure 6E). Therefore, the presence of the J-like domain of ARC6 enables it to modulate the inhibitory activity of ARC3 on FtsZ assembly.

**Figure 6.**
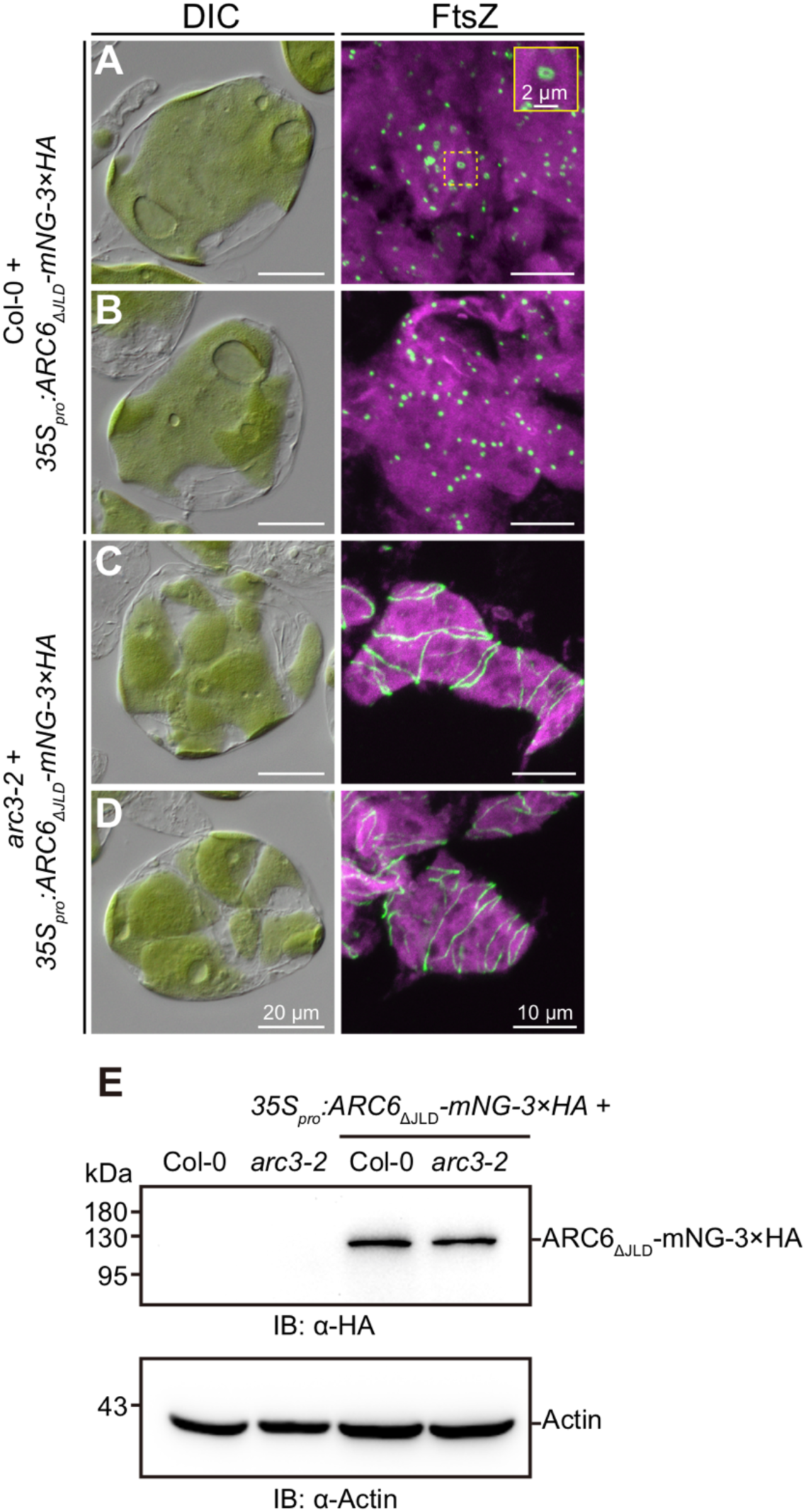
Overexpression of ARC6_ΔJLD_ causes disruption of Z-ring formation in an ARC3-dependent manner. **(A-D)** Chloroplast morphology (left panels) and FtsZ localization (right panels) in mesophyll cells of T_1_ transgenic plants expressing *35S_pro_:ARC6_ΔJLD_-mNeonGreen (mNG)-3×HA* in **(A, B)** wild-type Col-0 and **(C, D)** *arc3-2* mutant plants. Untransformed wild-type Col-0 and *arc3-2* mutant are shown in Figure 2, A and B. The inset in the right panel of **(A)** is a magnified image of the mini-ring, as indicated by the dashed line. Chloroplast morphology and FtsZ localization were observed using differential interference contrast (DIC) microscopy and immunofluorescence staining of FtsZ2-1, respectively. Scale bars are 20 μm for DIC images and 10 μm for all immunofluorescence images except the inset, which is 2 μm. **(E)** Immunoblot analysis of ARC6_ΔJLD_-mNG-3×HA from T_1_ transgenic Col-0 and *arc3-2* plants expressing *35S_pro_:ARC6_ΔJLD_-*mNG*-3×HA*. Total proteins extracted from leaf tissue of 4- to 5-week-old plants were separated on a 10% SDS-PAGE gel, and the membrane was probed with an anti-HA antibody. The expression of Actin, detected with an anti-Actin antibody, was used as a loading control.

### The J-like Domain of ARC6 Is Required for Chloroplast Division Activity but Not for ARC6 Localization

To further characterize the role of the J-like domain of ARC6 in chloroplast division, we generated constructs expressing *ARC6_ΔJLD_-mNG-3×HA*, *ARC6_ΔPPQ_-mNG-3×HA*, and *ARC6-mNG-3×HA*, all of which were driven by the *ARC6* native promoter. All transgenes were transformed back into the *arc6-5* mutant to evaluate their functionalities. As shown in Figure 7, ARC6-mNG-3×HA was able to restore both the chloroplast division defect and the disruption of Z-ring assembly in the *arc6-5* mutant (Figure 7, A and J), validating the functionality of this fusion protein *in vivo*. In contrast, neither ARC6_ΔJLD_-mNG-3×HA nor ARC6_ΔPPQ_-mNG-3×HA complemented the chloroplast morphology and Z-ring assembly phenotypes in the *arc6-5* transgenic plants (Figure 7, B to D). These differences were presumably not caused by the distinct expression levels of the fusion proteins since they were expressed at a similar level in the *arc6-5* transgenic plants (Figure 7I). Intriguingly, ARC6_ΔPPQ_ worked better than ARC6_ΔJLD_ in terms of complementing the division defects in the *arc6-5* transgenic plants (Figure 7, B to D, and J).

**Figure 7.**
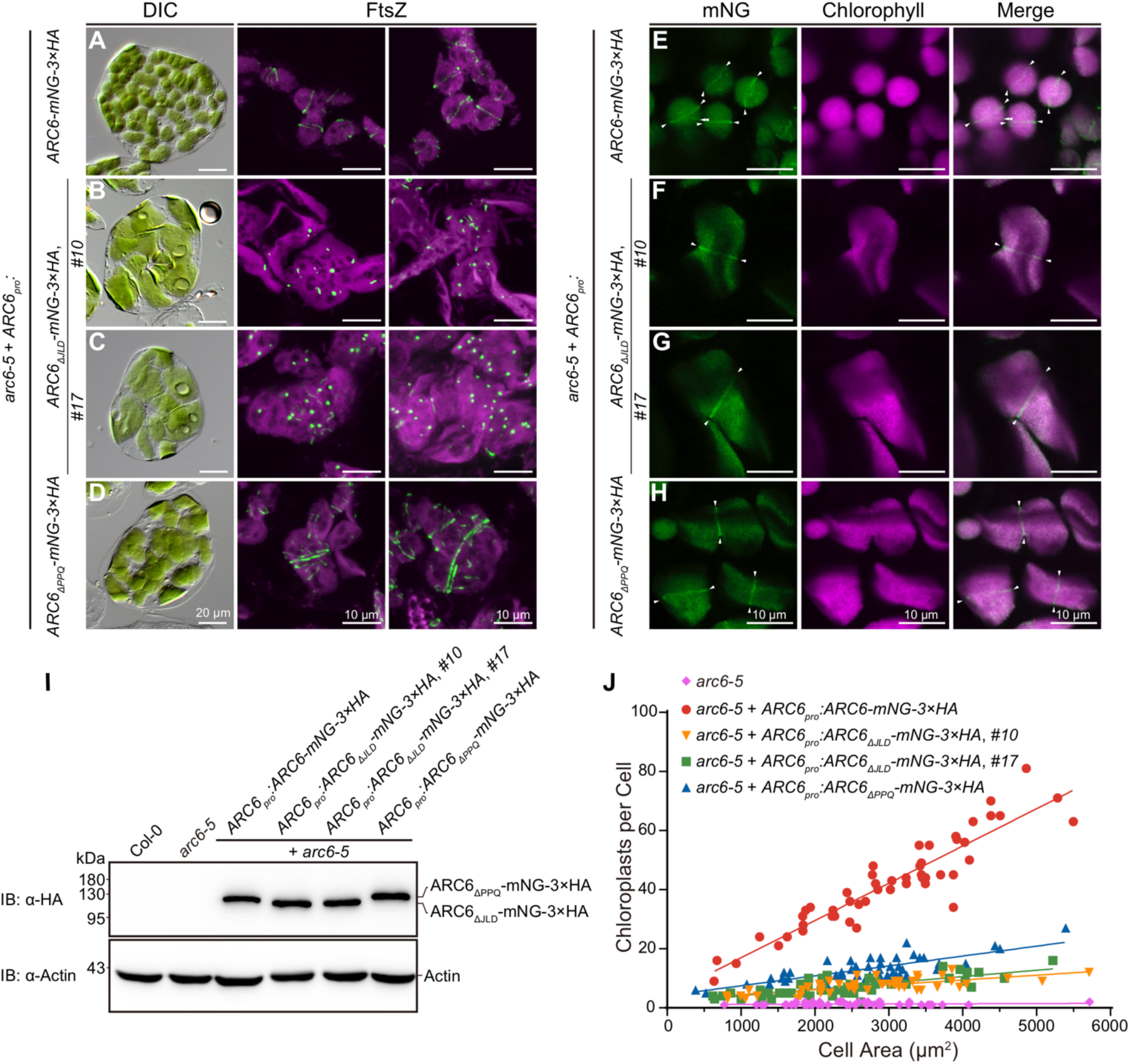
The J-like domain of ARC6 is required for chloroplast division but not for ARC6 localization. **(A-D)** Chloroplast morphology (left panels) and FtsZ localization (middle and right panels) in mesophyll cells of T_1_ transgenic plants expressing **(A)** *ARC6_pro_:ARC6-mNG-3×HA*, **(B, C)** *ARC6_pro_:ARC6_ΔJLD_-mNG-3×HA*, and **(D)** *ARC6_pro_:ARC6_ΔPPQ_-mNG-3×HA* in *arc6-5*, respectively. Untransformed *arc6-5* is shown in Figure 2C. Chloroplast morphology and FtsZ localization were observed using differential interference contrast (DIC) microscopy and immunofluorescence staining of FtsZ2-1, respectively. Scale bars are 20 μm for DIC images and 10 μm for all immunofluorescence images. **(E-H)** Subcellular localization of **(E)** ARC6-mNG-3×HA, **(F, G)** ARC6_ΔJLD_-mNG-3×HA, and **(H)** ARC6_ΔPPQ_-mNG-3×HA in 4- to 5-week-old T_1_ transgenic *arc6-5* plants. mNG signals (green) are shown in the left panels, and chlorophyll autofluorescence (magenta) is shown in middle panels. White arrowheads indicated ring-like structures formed by ARC6 and its derivative fusion proteins. Bars = 10 μm. (**I**) Immunoblot analysis of T_1_ transgenic *arc6-5* plants expressing *ARC6-mNG-3×HA*, *ARC6_ΔJLD_-mNG-3×HA*, and *ARC6_ΔPPQ_-mNG-3×HA*, respectively, under the control of the native *ARC6* promoter. Total proteins extracted from leaf tissue of 4- to 5-week-old plants were separated on a 10% SDS-PAGE gel, and the membrane was probed with an anti-HA antibody. The expression of Actin, detected with an anti-Actin antibody, was used as a loading control. (**J**) Quantitative analysis of chloroplast number versus mesophyll cell size (*n* = 50 cells) in the designated genotypes from **(A-D)**. The computed slopes of the best-fit lines are: *arc6-5*, 0.0001 (R^2^ = 0.015); *arc6-5* + *ARC6_pro_:ARC6-mNG-3×HA*, 0.0126 (R^2^ = 0.856); *arc6-5* + *ARC6_pro_:ARC6_ΔJLD_-mNG-3×HA* (#10), 0.0016 (R^2^ = 0.525); *arc6-5* + *ARC6_pro_:ARC6_ΔJLD_-mNG-3×HA* (#17), 0.0021 (R^2^ = 0.562); *arc6-5* + *ARC6_pro_:ARC6_ΔPPQ_-mNG-3×HA*, 0.0033 (R^2^ = 0.631).

To determine whether the J-like domain is required for ARC6 localization, we assessed the fluorescence signal of ARC6 fusion proteins in these transgenic plants. As previously reported (Vitha et al., 2003), we observed a midplastid ring-like structure for wild-type ARC6-mNG-3×HA (Figure 7E). In addition, we observed diffuse ARC6 signals in the chloroplasts of these plants (Figure 7E). To validate whether these signals derived from ARC6 fusion protein or autofluorescence from chlorophyll, we examined ARC6 in root tip and petal of flowers to exclude potential interference of chlorophyll autofluorescence. As in mesophyll cells, ARC6 exhibited diffuse localization in plastids from the root tip and petal cells (Supplemental Figure 9), suggesting that the diffuse signals found in the chloroplasts of the complemented plants were genuine ARC6. We found that both the ARC6_ΔJLD_ and ARC6_ΔPPQ_ fusion proteins formed ring-like structures in addition to being diffuse in chloroplasts from transgenic *arc6-5* plants (Figure 7, F to H). These data demonstrate that the J-like domain of ARC6 is required for its activity but not for its localization during chloroplast division.

### ARC6 Recruits ARC3 to the Chloroplast Division Site

In a previous study (Chen et al., 2019), we demonstrated that ARC3 localizes to two distinct pools in the chloroplast: one is diffusely distributed throughout the stroma, and the other forms a ring-like structure at the division site. Since ARC6 localizes at the midplastid division site and interacts with ARC3, we wondered whether it directly recruits ARC3 to form the ring-like structure at the division site. Towards addressing this, we employed a construct encoding an ARC3-mNeonGreen (ARC3-mNG) fusion protein driven by its native promoter, whose functionality has been previously validated (Figure 8A) (Chen et al., 2019). We transformed this construct into the *arc3-2 arc6-5* double mutant and observed the distribution of ARC3-mNG in the transgenic plants. As previously reported (Chen et al., 2019), ARC3-mNG formed a ring-like structure at the middle of the chloroplast (Figure 8, A to C), in addition to being diffusely distributed throughout the chloroplast stroma (Figure 8, A to C). In contrast, we barely observed any ARC3-mNG ring structures in the *arc3-2 arc6-5* transgenic mutant (Figure 8, D to F, M). However, the pool of diffuse ARC3-mNG still existed despite the absence of ARC6 (Figure 8, D to F), indicating that localization of the diffuse ARC3 is not regulated by ARC6. Statistical analysis showed that the percentage of ARC3-mNG ring structures formed in the absence of ARC6 (10.1%) decreased significantly compared to the percentage in its presence (53.8%) (Figure 8M). Western blot analysis confirmed that ARC3-mNG was expressed at a similar level in the complemented *arc3-2* transgenic line and *arc3-2 arc6-5* transgenic line (Figure 8N).

**Figure 8.**
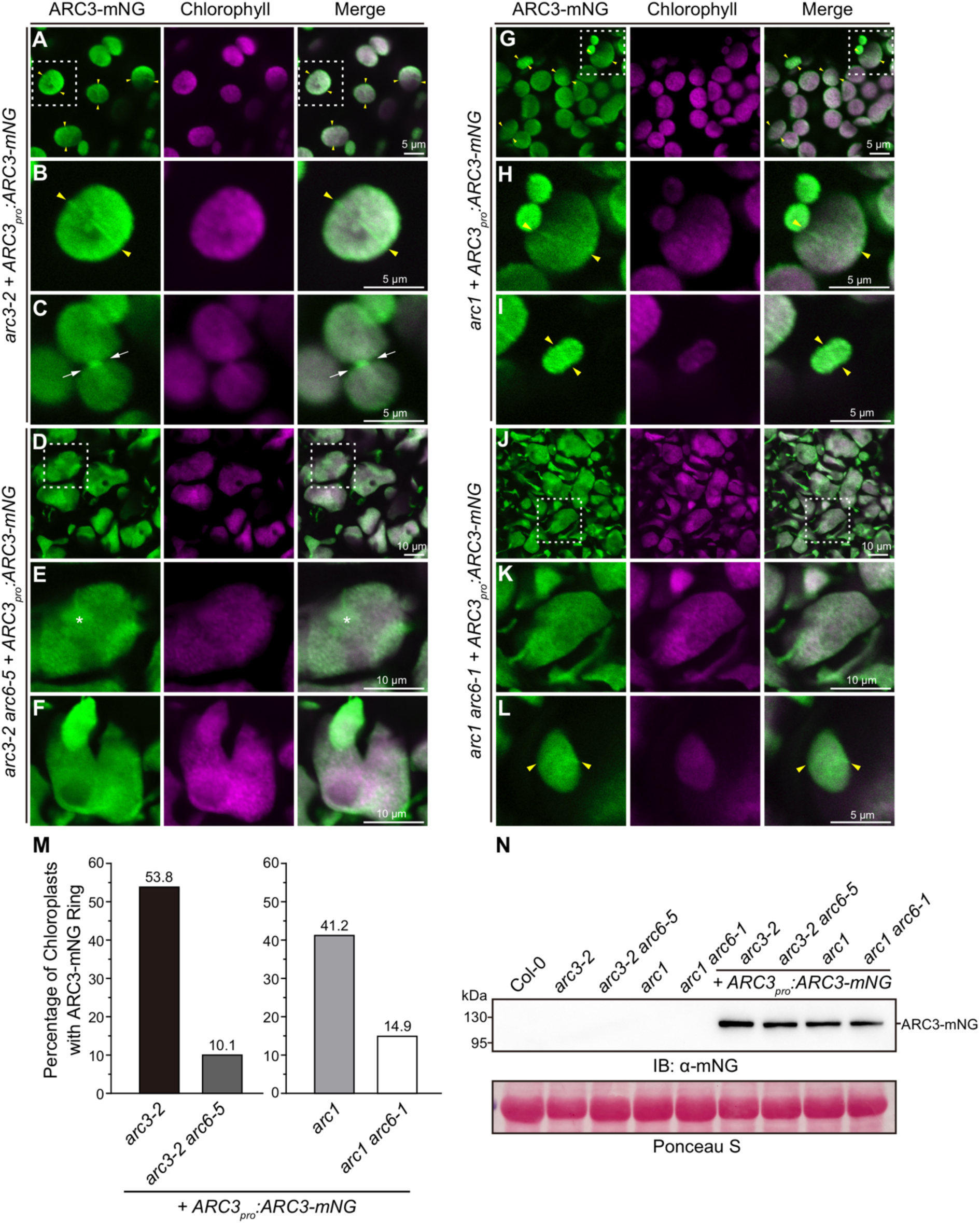
ARC6 recruits ARC3 to the chloroplast division site. **(A-L)** Subcellular localization of ARC3-mNG in transgenic **(A-C)** *arc3-2*, **(D-F)** *arc3-2 arc6-5*, **(G-I)** *arc1*, and **(J-L)** *arc1 arc6-1* mutant plants expressing *ARC3_pro_:ARC3-mNG*. The mNG fluorescence (green) and chlorophyll autofluorescence (magenta) signals were acquired through confocal laser scanning microscopy. Midplastid-localized ARC3-mNG ring structures are associated with both **(A, B, G-I, L)** unconstricted chloroplasts (yellow arrowheads) and **(C)** constricted chloroplasts (white arrows). The white double-arrowheads in **(A)** indicate the presence of additional ARC3-mNG strands that are not affiliated with the division site. Asterisk indicates an ARC3-mNG filament instead of an intact ring in a giant chloroplast of transgenic *arc3-2 arc6-5*. The regions enclosed by the white dashed box in **(A, D, G, J)** are magnified in **(B, E, H, K)**. Bars are as indicated. **(M)** Quantification of ARC3-mNG ring structures in chloroplasts of transgenic arc3*-2*, *arc3-2 arc6-5*, *arc1*, and *arc1 arc6-1* plants expressing *ARC3_pro_:ARC3-mNG*. The percentage of chloroplasts with ARC3-mNG rings was 53.8% (*n* = 145) in *arc3-2*, 10.1% (*n* = 158) in a*rc3-2 arc6-5*, 41.2% (*n* = 143) in *arc1* and 14.9% (*n* = 161) in *arc1 arc6-1*. *n* is the total number of chloroplasts analyzed. **(N)** Immunoblot analysis of ARC3-mNG fusion protein in transgenic *arc3-2*, a*rc3-2 arc6-5*, *arc1*, and *arc1 arc6-1* plants expressing *ARC3_pro_:ARC3-mNG*. Total proteins extracted from leaf tissue of 4-week-old plants were separated on a 10% SDS-PAGE gel, and membrane was probed with an anti-mNG antibody. Ponceau S-stained large subunit of Rubisco (bottom panel) was served as a loading control.

In the *arc6-5* mutant, there are only 1∼2 chloroplasts with dramatically enlarged size (Figure 2D). To exclude the possibility that the lack of ARC3-mNG ring formation was due to the giant chloroplast size in the *arc6-5* mutant, we employed the *arc1 arc6-1* double mutant. ARC1 (also known as FtsHi1) is a negative regulator of chloroplast division, and the division defect is partially restored in the *arc1 arc6* double mutant (Supplemental Figure 10) (Kadirjan-Kalbach et al., 2012). We therefore transformed the *ARC3-mNG* construct into the *arc1* and *arc1 arc6-1* mutants (Figure, G to L). As in the complemented *arc3-2* transgenic line (Figure 8, A to C), the ARC3-mNG ring was observed in the transformed *arc1* plants (Figure 8, G to I). We noticed that the percentage of ARC3-mNG ring was lower in *arc1* (41.2%) than in *arc3-2* (53.8%) transgenic plants (Figure 8M), which could be, at least partially, due to the endogenous expression of non-tagged ARC3 in the *arc1* mutant. Our results showed that the percentage of ARC3-mNG rings detected in *arc1 arc6-1* (14.9%) was much lower than in the *arc1* transgenic plants (41.2%) (Figure 8M). The expression level of the ARC3-mNG fusion protein was similar among the distinct background plants (Figure 8N). Taken together, these data demonstrate that ARC6 recruits ARC3 to the midplastid during chloroplast division.

## DISCUSSION

ARC6 is considered one of the central components of the chloroplast division machinery (Pyke et al., 1994; Vitha et al., 2003; Maple et al., 2005; Glynn et al., 2008; Johnson et al., 2013; Wang et al., 2017). The molecular mechanisms underlying ARC6’s function in chloroplast division, however, remain largely unknown. In this study, ARC3 was identified as a novel interactor of ARC6, and the biological significance of the ARC6-ARC3 interaction was investigated. Our work provided compelling evidence that ARC6 regulates the formation and positioning of the Z ring by fine-tuning ARC3 activity at the chloroplast division site. A working model depicting the function of the ARC6-ARC3 complex at the midplastid site is presented in Figure 9.

**Figure 9.**
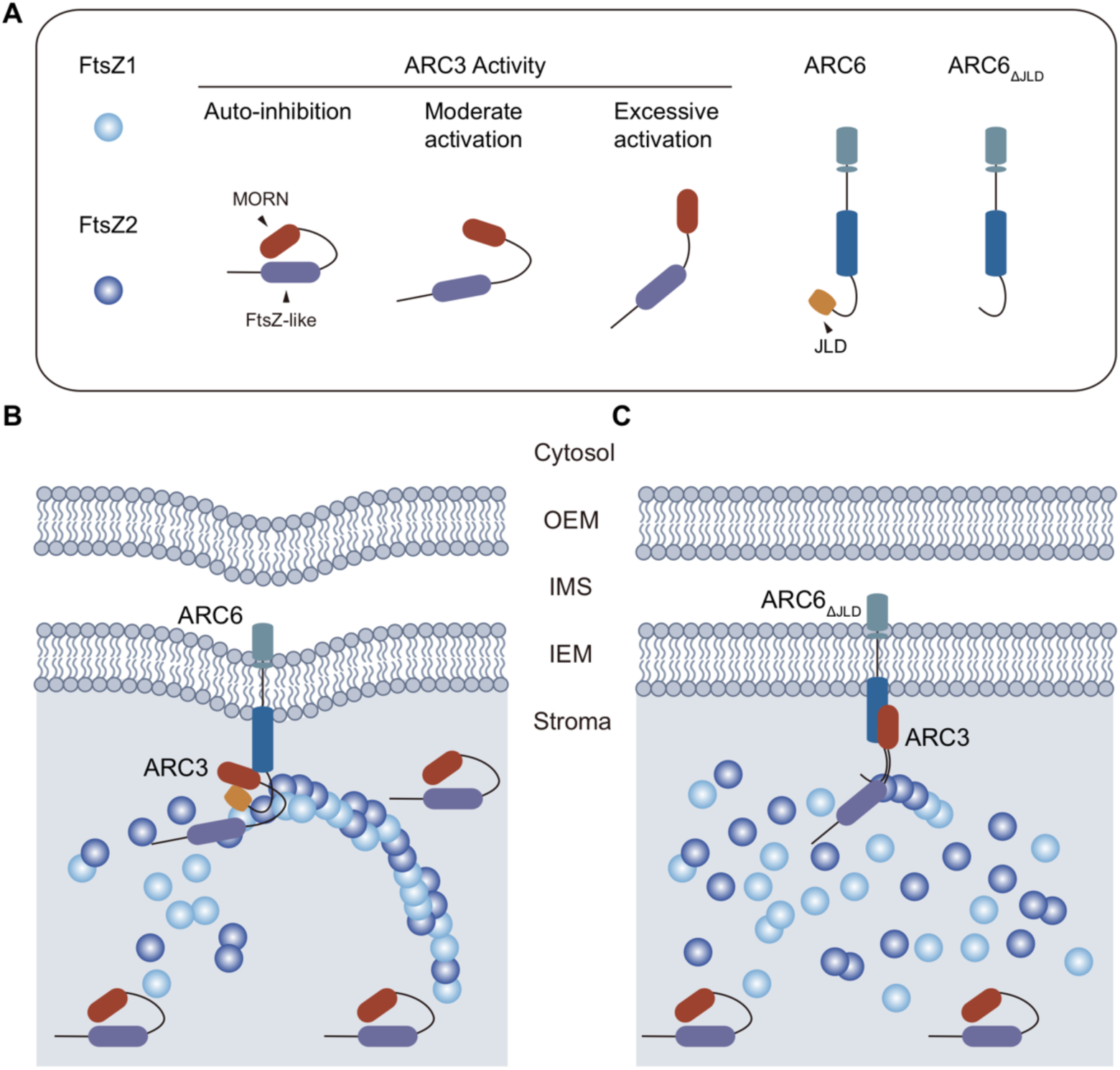
Working model of the ARC6-ARC3 complex in the regulation of Z-ring assembly during chloroplast division. (A) Full-length ARC3 is inhibited from interacting with FtsZ proteins due to the presence of the MORN domain and thus is inactive in terms of inhibiting Z-ring assembly or accelerating Z-ring dynamics during the constriction of chloroplasts (Zhang et al., 2013; Chen et al., 2019). Binding of ARC3 by wild-type ARC6 activates ARC3 to a moderate extent. ARC6_ΔJLD_, the JLD-deleted version of ARC6, however, binds ARC3 more strongly and thus results in overactivated ARC3. (B) The FtsZ-ring, comprising heteropolymers of FtsZ1 and FtsZ2 (Vitha et al., 2001; Yoder et al., 2007; Yoshida et al., 2016), anchors to the chloroplast inner membrane through the interaction between FtsZ2 and ARC6 (Maple et al., 2005; Johnson et al., 2013). ARC6 recruits ARC3 to the chloroplast division site and activates the inhibitory activity of ARC3 on the assembly of FtsZ filaments. Such activation may result from the binding of the ARC3 MORN domain by the JLD of ARC6, which may alter the conformation of ARC3. Binding of ARC3 by ARC6 leads to moderately activated ARC3, which is able to inhibit non-specific Z-ring formation in the vicinity of the division site and accelerate Z-ring dynamics during constriction of chloroplasts (Chen et al., 2019). (C) ARC6_ΔJLD_, the JLD-deleted version of ARC6, binds and activates full-length ARC3 more strongly than ARC6. Overexpression of ARC6_ΔJLD_ leads to excessive activation of ARC3, which causes disruption of the assembly and formation of the Z ring. MORN, Membrane Occupation and Recognition Nexus domain; JLD, J-like domain; OEM, outer envelope membrane; IMS, intermembrane space region; IEM, inner envelope membrane.

ARC6 and ARC3 were found to be co-precipitated within a protein complex obtained from Arabidopsis (McAndrew et al., 2008), suggesting a potential direct interaction between them during chloroplast division. We investigated the ARC6-ARC3 protein interaction in the current study using yeast and plant systems. In contrast to a previous report (Maple et al., 2007), our Y2H result showed a positive interaction between ARC6 and ARC3. The lack of such interaction in prior study could be due to the inclusion of transit peptides in their Y2H assay, which could interfere with the interaction between the two proteins (Maple et al., 2007). Transit peptides of the precursor chloroplast proteins are removed after import into the chloroplast and thus are not part of the mature chloroplast proteins (Bruce, 2000; Lee and Hwang, 2018; McKinnon and Theg, 2019). In the current study, the predicted transit peptides of both proteins were excluded when performing this assay. Our Co-IP and split-LUC assays further validated the ARC6-ARC3 interaction *in planta*. In contrast, the prior study failed to detect such an interaction using the bimolecular fluorescence complementation (BiFC) assay in tobacco plants (Maple et al., 2007). We revisited the detailed methods for the BiFC experiments and noticed that the fluorescent tag was fused to the C-terminus of ARC6, which faces the intermembrane space of the chloroplast instead of the stroma where ARC3 localizes (Vitha et al., 2003; Maple et al., 2007). We therefore considered the absence of the ARC6-ARC3 interaction reported by Maple et al. (2007) a false negative. Thus, we identified ARC3 as a novel interactor of ARC6.

ARC6 is known to be a positive regulator of Z-ring assembly, as the *arc6* mutant shows only short FtsZ fragments or patches (Vitha et al., 2003). A recent *in vitro* study suggested that this may be achieved through ARC6’s prevention of the disassembly of GDP-bound FtsZ proteins from the assembled filaments, resulting in the stabilization of the assembled FtsZ filaments (Sung et al., 2018). In contrast, we showed that ARC6 not only interacts with ARC3, the direct inhibitor of Z-ring assembly (Zhang et al., 2013), but also activates the inhibitory activity of ARC3 on FtsZ filament assembly, which presumably leads to the inhibition of Z-ring assembly by ARC3 *in vivo*. This is the first report of these novel functions of ARC6, highlighting the complexity of ARC6’s role during chloroplast division. Given that ARC6 binds to FtsZ2 as well (Maple et al., 2005; Schmitz et al., 2009), it’s difficult to test the effect of the ARC6-ARC3 complex on the dynamics of the reconstituted Z ring in the heterologous yeast *Pichia pastoris*, as we did for the PARC6-ARC3 complex previously (Chen et al., 2019). However, we previously showed that the mid-plastid localized ARC3 functions to prevent non-specific Z-ring formation in the vicinity of the division site as well as to promote the dynamics of the Z ring in order to facilitate Z-ring constriction and completion of chloroplast division (Chen et al., 2019). Thus, we propose that activation of ARC3 by ARC6 functions similarly at the division site. In addition, ARC6, ARC3, and FtsZ2 may form a complex to regulate chloroplast division. This was initially evidenced by the finding that ARC6 and ARC3 co-existed in a native FtsZ complex purified from the leaf of the Arabidopsis plant (McAndrew et al., 2008). In accordance with this, our Y3H data validated the formation of the ARC6-ARC3-FtsZ2 complex since the transformed cell can grow on the more stringent –His –Ade selective plates. In contrast, the transformed cells expressing ARC6, ARC3, and FtsZ1 cannot grow on such stringent selective plates since they lack a direct interaction between ARC6 and FtsZ1. Overall, our results demonstrate that ARC6 functions together with ARC3 and FtsZ2 during chloroplast division.

The establishment and subsequent constriction of the chloroplast division machinery require coordination of the stromal Z ring with the cytosolic DRP5B ring across the chloroplast membranes (Osteryoung and Pyke, 2014; Chen et al., 2018a). As an inner envelope membrane protein, ARC6 is a key player in such coordinated behavior. The stromal region of ARC6 interacts with FtsZ2 and is mainly responsible for tethering the stromal localized Z ring to the inner chloroplast envelope membrane (Maple et al., 2005; Johnson et al., 2013). The intermembrane space (IMS) region of ARC6 interacts with the outer envelope membrane protein PDV2 and recruits it to the chloroplast division site (Glynn et al., 2008). Recently, PDV2 was shown to induce the dimerization of ARC6 through interaction in the IMS region of the chloroplast (Wang et al., 2017). The lack of such dimerization in the *pdv2* mutant leads to mislocalization of ARC6 at the chloroplast division site and results in clustered ARC6 ring-like structures around the middle of the chloroplasts in the *pdv2* mutant (Wang et al., 2017). The positioning of the Z ring is ultimately governed by ARC3, in the context of the chloroplast Min system (Maple et al., 2007; Nakanishi et al., 2009; Zhang et al., 2013). However, it has been shown that there are multiple Z rings in the *pdv2* mutant, suggesting that ARC3 is not fully functional in the absence of PDV2 (Miyagishima et al., 2006). Thus, we hypothesize that the presence of PDV2 may be required for stable ARC6-ARC3 interaction at the chloroplast division site. Further study is required to investigate the ARC6-ARC3 interaction in the *pdv2* mutant and its impact on chloroplast division.

ARC6 is a non-canonical J protein since the J-like domain within ARC6 lacks the conserved tripeptide HPD (Vitha et al., 2003; Pulido and Leister, 2018). Consistent with this, we failed to detect a direct interaction between the J-like domain of ARC6 and the chloroplastic Hsp70 proteins (Supplemental Figure 3). Thus, ARC6 was designated as a DNAJD protein in recent literatures since it contains only a J-like domain with an unknown function (Pulido and Leister, 2018; Tamadaddi et al., 2022). The Y2H assays performed with ARC6 derivates, missing the intact J-like domain or only the conserved PPQ motif, suggested that the J-like domain of ARC6 functions to reduce the binding strength between ARC6 and ARC3. This was further supported by the data obtained from the heterologous yeast *S. pombe*, where the ARC6_ΔJLD_ activated the inhibitory activity of ARC3 much more than full-length ARC6. Moreover, overexpression of ARC6_ΔJLD_ in wild-type Col-0 led to the disruption of Z-ring assembly and formation, resembling the phenotypes observed in ARC3 overexpression transgenic plants (Zhang et al., 2013). The ability of ARC6_ΔJLD_ to do so, however, depended on the availability of ARC3, since no such disruption of the Z ring was detected when ARC6_ΔJLD_ was expressed at a similar level in *arc3-2* mutant as compared to the wild-type Col-0 plant, implying that it acts through activating ARC3. Given that ARC6_ΔPPQ_ behaved like ARC6_ΔJLD_ in terms of promoting the interaction between full-length ARC3 and the FtsZ proteins, we proposed that overexpression of ARC6_ΔPPQ_ would cause the disruption of Z-ring formation in an ARC3-dependent manner as well. Our data suggested that the major function of the J-like domain of ARC6 is to fine-tune the activity of ARC3 to keep it at a proper level at the chloroplast division site. By doing so, ARC3 could be active enough to prevent the non-specific assembly of Z rings in the vicinity of the division site but not be too active to inhibit Z-ring assembly and formation (Chen et al., 2019). Interestingly, we noticed that ARC6_ΔPPQ_ was more efficient than ARC6_ΔJLD_ in complement of the chloroplast division defect in the *arc6* mutant. Our Y2H data showed that the interaction between ARC6 and FtsZ2 was reduced by the depletion of the J-like domain while not affected by the removal of only the PPQ motif. Thus, the functional difference between ARC6_ΔPPQ_ and ARC6_ΔJLD_ in chloroplast division could be attributed, at least partially, to the distinct binding affinity of these ARC6 derivatives with FtsZ2. Our findings demonstrated that the biological function of the ARC6 J-like domain is to modulate the binding intensity of the ARC6-ARC3 interaction and the subsequent activation of ARC3 by ARC6, as well as to promote the ARC6-FtsZ2 interaction.

We previously reported that PARC6 activates ARC3 to promote Z-ring dynamics at the chloroplast division site (Chen et al., 2019). Moreover, we noticed that PARC6 partially activates ARC3 (Zhang et al., 2016; Chen et al., 2019). Interestingly, it seems that ARC6 partially activates ARC3 as well in the current study. It’s unclear whether PARC6 and ARC6 will interfere with each other in terms of regulating the activity of ARC3 at the chloroplast division site. Regarding the regulation of ARC3 activity, ARC6 and PARC6 presumably work cooperatively to maintain the inhibitory activity of ARC3 at the proper level at the chloroplast division site. Thus, we propose that the activity of ARC3 is under strict control by multiple factors to ensure both the correct positioning and the proper dynamics of the Z ring during chloroplast division. Given the morphology of the Z ring in the *arc6* mutant (Vitha et al., 2003), one possible function of ARC6 is to inhibit the activity of the chloroplast Min system. Since ARC3 is the core component of the Min system, the J-like domain is likely to regulate the interaction between ARC6 and other Min components as well. It has been reported that ARC6 interacts with both MinD and MCD1 (Chen et al., 2018b; Zhang et al., 2021). It would be interesting to investigate whether the J-like domain could regulate the interaction between ARC6 and these Min system components in future studies.

## METHODS

### Plant Materials and Growth Conditions

*Arabidopsis thaliana* ecotype Columbia-0 (Col-0) and Landsberg *erecta* (L*er*) were used as the wild type in this study. The *arc3-2* (SALK_057144) and *arc6-5* (SAIL_693_G04) are in the Col-0 background (Shimada et al., 2004; Glynn et al., 2008); *arc1* is in the L*er* background, and *arc6-1* is in the Wassilewskija (WS) background (Pyke and Leech, 1992; Pyke et al., 1994; Vitha et al., 2003; Kadirjan-Kalbach et al., 2012). The *arc3-2 arc6-5* double mutant was generated through genetic crossing and confirmed by genotyping. All seeds were surface-sterilized in 70% (v/v) ethanol with 0.05% (v/v) Triton X-100 and sown on half-strength Murashige and Skoog Basal Medium with Vitamins (PhytoTech, M519) in 1% (w/v) sucrose and 0.7 (w/v) agar at pH 5.7. The seeds were kept in the dark for 2 d at 4°C and then transferred to the growth room (100 μmol m^-2^ s^-1^) at 22°C with a 16-h light/8-h dark cycle and 70% humidity. *Nicotiana benthamiana* plants were grown (125 μmol m^-2^ s^-1^) at 23°C with the identical photoperiod and humidity of Arabidopsis.

### Plasmid Design and Construction Strategies

All chloroplast division genes expressed in yeast cells were amplified from the cDNA sequences of *Arabidopsis thaliana* and did not contain the coding sequences for the predicted transit peptides. Regarding expression *in planta*, the chloroplast division genes were amplified from the genomic sequences. All PCR amplifications were performed using Phanta Max Super-Fidelity DNA Polymerase (Vazyme, P505) in a Biometra Tone 96 G, 230 V Thermocycler (Analytik Jena). Restriction enzymes were acquired from New England Biolabs and TransGen Biotech. The Gibson Assembly approach was employed for the construction of all vectors (Gibson et al., 2009). The primers are detailed in Supplemental Table 1, and the obtained constructs were verified by sequencing prior to further use.

The pGADT7 (AD) and pGBKT7 (BD) vectors (Clontech) served as the backbone for generating constructs in the yeast two-hybrid (Y2H) and yeast three-hybrid (Y3H) assays. The CC268/CC269 primer set was used to amplify the *ARC6_68-614_* coding sequence, and the obtained fragment was inserted into the AD vector digested with NdeI and BamHI, yielding pGADT7-ARC6_68-614_. The CC268/CC270 and CC271/CC269 primer sets amplified *ARC6_ΔJLD_* from pGADT7-ARC6_68-614_, and the resulting fragment was cloned into the AD vector digested with NdeI and BamHI, yielding pGADT7-ARC6_ΔJLD_. The CC268/WB352 and WB353/WB354 primer sets were used to amplify *ARC6_ΔPPQ_* from pGADT7-ARC6_68-614_, and the product was inserted into the AD vector digested with NdeI, yielding pGADT7-ARC6_ΔPPQ_. The rest of the constructs employed in the Y2H assays were described previously (Glynn et al., 2009; Zhang et al., 2013; Zhang et al., 2016; Chen et al., 2019).

To generate constructs used in the Y3H assays, the *ARC6_68-614_* expression cassette (without the GAL4 activation domain) was amplified from pGADT7-ARC6_68-614_ with CC172/CC169 and CC170/CC173 primers. The obtained fragments were inserted into AvrII-digested pGBKT7-FtsZ1, pGBKT7-FtsZ2 and pGBKT7-Empty, respectively, to generate pGBKT7-FtsZ1; ARC6, pGBKT7-FtsZ2; ARC6 and pGBKT7-Empty; ARC6 vectors. Similarly, the *ARC6_ΔJLD_* and the *ARC6_ΔPPQ_* expression cassettes, both deprived of the GAL4 activation domain, were amplified from pGADT7-ARC6_ΔJLD_ and pGADT7-ARC6_ΔPPQ_, respectively, with the identical CC172/CC169 and CC170/CC173 primer sets. The obtained fragments were inserted into AvrII-digested pGBKT7-FtsZ1, pGBKT7-FtsZ2 and pGBKT7-Empty, respectively, in order to obtain the rest of Y3H constructs.

The pGREEN and pCAMBIA1300-20 (Zhang et al., 2013) vectors were used as the backbone to generate constructs for the Co-IP assays. The genomic sequence of *ARC3* was amplified with WB378/WB379 primer set, and the obtained product was cloned into XhoI/PstI digested pGREEN, leading to pGREEN-35S_pro_:ARC3-6×HA. Likely, the genomic sequence of *FtsZ2* was amplified with WB382/WB383 primers, and the fragment was inserted into pGREEN vector digested with XhoI and PstI, in order to make pGREEN-35S_pro_:FtsZ2-6×HA. To generate pCAMBIA1300-20-35S_pro_:ARC6-eGFP, the sequences for *ARC6* (genomic) and *eGFP* as well as *3’UTR* of *ARC6* were amplified using primer set WB345/WB339, WB340/WB341 and WB342/WB343, respectively. The obtained three fragments were inserted into pCAMBIA1300-20 vector digested with BamHI and SacI through Gibson Assembly, resulting in the pCAMBIA1300-20-35S_pro_:ARC6-eGFP construct. The transit peptide of RecA (RecA_TP_) has been shown to target fusion proteins into chloroplasts (Kohler et al., 1997; Chen et al., 2018b). The pCAMBIA1300-20-35S_pro_:RecA_TP_-eGFP construct was generated by amplifying *RecA_TP_* and *eGFP* sequences using primer set WB69/WB497 and WB340/WB498, respectively. Subsequently, the resulting fragments were then inserted into pCAMBIA1300-20 vector digested with BamHI and SacI.

The pCAMBIA1300 vector was used as the backbone to generate constructs for the split-luciferase complementation (split-LUC) assays. To amplify *N-terminal luciferase* (*NLuc*), *C-terminal luciferase* (*CLuc*), and *6×HA* fragments, we used pCAMBIA1300-NLuc, pCAMBIA1300-CLuc, and pGREEN-35Spro:6×HA as templates. To generate pCAMBIA1300-ARC6_TP_-NLuc-ARC6, the primer sets WB460/WB461, WB462/WB463, WB464/WB465 and WB466/WB467 (for amplifying *6×HA* sequence) were used. The obtained fragments were then cloned into pCAMBIA1300 digested with KpnI and PstI. To make the pCAMBIA1300-ARC3-CLuc and pCAMBIA1300-FtsZ2-CLuc constructs, the pCAMBIA1300 backbone was digested with SacI and SalI. The primer sets WB468/WB469, WB470/WB471 and WB472/WB473 were used to produce fragments for *ARC3-CLuc* transgene; WB474/WB475, WB470/WB471 and WB472/WB473 were used to generate fragments for *FtsZ2-CLuc* transgene. The sequencing results indicated that a 3×HA, instead of 6×HA, was fused with FtsZ2-CLuc.

The pREP41X and pREP42X vectors (http://www-bcf.usc.edu/; forsburg/vectortable.html) were employed as the backbone for constructing plasmids expressing chloroplast division proteins in yeast *Schizosaccharomyces pombe* (*S. pombe*). The *ARC6_68-614_* and *ARC6_ΔJLD_* fragments were amplified with the primers CC101/CC102. The *mRuby2* fragment was amplified using the CC85/CC86 primers from pREP41X-mRuby2. These fragments were then cloned into BamHI-digested pREP41X to generate pREP41X-ARC6_68-614_-mRuby2 and pREP41X-ARC6_ΔJLD_-mRuby2 constructs, respectively. To generate pREP41X-ARC3_41-741_-mCerulean; ARC6_68-614_-mRuby2 and pREP41X-ARC3_41-741_-mCerulean; ARC6_ΔJLD_-mRuby2, primers AT109/AT110 were used to amplify the *ARC6_68-614_-mRuby2* and *ARC6_ΔJLD_-mRuby2* expression cassettes (including the *nmt1\** promoter and terminator sequences) from pREP41X-ARC6_68-614_-mRuby2 and pREP41X-ARC6_ΔJLD_-mRuby2, respectively. The obtained PCR products were inserted into AatII-digested pREP41X-ARC3_41-741_-mCerulean, yielding pREP41X-ARC3_41-741_-mCerulean; ARC6_68-614_-mRuby2 and pREP41X-ARC3_41-741_-mCerulean; ARC6_ΔJLD_-mRuby2 constructs, respectively. Likewise, the AT109/AT110 primer set was used to amplify the *mRuby2* expression cassette (including the *nmt1\** promoter and terminator sequences) from pREP41X-mRuby2, followed by insertion into pREP41X-ARC3_41-598_-mCerulean digested with AatII, generating pREP41X-ARC3_41-598_-mCerulean; mRuby2 construct as a control. The rest of constructs used in *S. pombe*, including pREP41X-ARC3_41-741_-mCerulean, pREP41X-ARC3_41-598_-mCerulean, pREP42X-FtsZ1-mVenus and pREP41X-ARC3_41-741_-mCerulean; mRuby2 were described in previous reports (TerBush et al., 2016; TerBush et al., 2018; Chen et al., 2019).

The pCAMBIA1300-20 vector was used as the backbone to generate constructs expressing distinct ARC6 derivates and ARC3 in Arabidopsis plants. To generate construct expressing *ARC6_pro_:ARC6(g)-mNeonGreen* (*mNG*)*-3×HA*, primers WB336/WB337 and WB338/WB339 were used to amplify a fragment harboring the promoter region of *ARC6* (∼2 kb) and *ARC6* genomic sequence; primers WB520/WB521, WB522/WB523 and WB342/WB343 were used to amplify *mNG*, *3×HA*, and the *3’UTR* (276 bp) of *ARC6*. Subsequently, the resulting fragments were inserted into pCAMBIA1300-20 backbone after digestion with PstI and SacI. To obtain construct expressing *ARC6_pro_:ARC6_ΔJLD_(g)-mNG-3×HA*, primers WB336/WB337 and WB344/WB339 were used to amplify a fragment containing the promoter region of *ARC6* (∼2 kb) and *ARC6_ΔJLD_* genomic sequence. Likewise, primers WB336/WB352 and WB353/WB339 were used to amplify a fragment containing the promoter region of *ARC6* (∼2 kb) and *ARC6_ΔPPQ_* genomic sequence in order to make *ARC6_pro_:ARC6_ΔPPQ_(g)-mNG-3×HA*. Regarding the identical part of the two transgenes, primers WB520/WB521, WB522/WB523 and WB342/WB343 were used to amplify *mNG*, *3×HA*, and the *3’UTR* (276 bp) of *ARC6*. The obtained fragments were then cloned into PstI/SacI-digested pCAMBIA1300-20 to generate the destinated constructs expressing *ARC6_pro_:ARC6_ΔJLD_(g)-mNG-3×HA* and *ARC6_pro_:ARC6_ΔPPQ_(g)-mNG-3×HA*. To obtain *35S_pro_:ARC6(g)-mNG-3×HA*, primers of WB345/WB339, WB520/WB523 and WB342/WB343 were used to amplify fragments of ARC6 genomic region, *mNG-3×HA* and *3’ UTR* of *ARC6*. To make *35S_pro_:ARC6_ΔJLD_ (g)-mNG-3×HA*, primers WB345/WB337, WB338/WB339, WB520/WB523 and WB342/WB343 were used to amplify fragments of *ARC6_ΔJLD_* genomic region, *mNG-3×HA* and *3’ UTR* of *ARC6*. These fragments were inserted into the pCAMBIA1300-20 digested with BamHI and SacI in order to generate the final constructs. Construction of *ARC3_pro_:ARC3-mNG* was described previously (Chen et al., 2019).

### Generation of Transgenic Plants

All transgenic plants were generated by floral dipping (Clough and Bent, 1998) using the *Agrobacterium tumefaciens* GV3101 strain. Transgenic plants were screened on half-strength Murashige and Skoog Basal Medium (PhytoTech, M519) supplemented with 20 μg/mL hygromycin (Yeasen, 60225ES03). After one week of growth on selective plates, positive transgenic seedlings were transferred to soil and cultivated in the growth room. T_1_ plants were utilized for all transgenic backgrounds, except for transgenic *arc3-2* expressing *ARC3_pro_:ARC3(g)-mNG*, where T_3_ plants were used.

### Chloroplast Phenotype Analysis

For the analysis of chloroplast phenotypes, young expanding leaves from 3-week-old plants were fixed according to previously established protocol (Pyke and Leech, 1991). Briefly, leaf tissue samples were fixed in 3.5% (v/v) glutaraldehyde for 1 hour, followed by transfer into 0.1 M Na_2_EDTA (pH = 9.0). The samples were kept at 50°C for 2 h, and were stored overnight at 4°C before observation. Mesophyll cells were examined using differential interference contrast optics equipped on an epifluorescence microscope (Olympus BX53) and imaged with an Olympus DP23 camera. The cell plane area was quantified using Fiji (ImageJ) software (http://fiji.sc/Fiji), and the number of chloroplasts in individual cells was counted manually.

### Immunofluorescence Staining of FtsZ Localization

FtsZ assembly and Z-ring positioning were analyzed by immunofluorescence staining utilizing an anti-FtsZ2-1 antibody as described previously (Vitha et al., 2001; Yoder et al., 2007; Vitha and Osteryoung, 2011; Chen et al., 2019). Briefly, young expanding rosette leaves were collected from 4- to 5-week-old plants and then cut into 2 mm width strips. The samples were embedded with wax (PEG distearate, Sigma-Aldrich 305413; 1-Hexadecanol, Sigma-Aldrich 258741). The sections were cut to a thickness of 5 μm using a Manual Rotary Microtome (Thermo Fisher Scientific). Sections were incubated with the primary anti-FtsZ2-1 antibody (1:3500 dilution), followed by incubation with the Alexa Fluor 488 Goat Anti-Rabbit secondary antibody (1:500 dilution) (Yeasen, 33106ES60). Images shown in Figure 6 and 7, and Supplemental Figure 7 and 8 were acquired with a Nikon A1 HD25 laser scanning confocal microscope. The confocal channel settings for detecting the FtsZ2-1 signal were 488 nm excitation with emission recorded between 500 and 550 nm, while chlorophyll autofluorescence was captured using 561 nm excitation and emission recorded between 662 and 737 nm. Single images were stacked using a maximum intensity projection method after Z-axis scanning in the confocal microscope. Images were merged by Fiji (ImageJ) software (http://fiji.sc/Fiji). Samples in Figure 2 were observed using an Olympus FV3000 laser scanning confocal microscope, and samples in Supplemental Figure 10 were observed with a Zeiss LSM-900 laser scanning confocal microscope. The parameters for the channel setting were similar to those mentioned for the Nikon A1 confocal.

### Yeast Two-Hybrid and Yeast Three-Hybrid Assays

The *HIS3* and *ADE2* reporter genes were employed to assess the protein interactions in the yeast strain Y2HGold (Takara Bio). Yeast transformation was carried out using the lithium acetate (LiAc)-mediated method. Positive transformants were selected based on growth on synthetic dropout (SD) medium lacking leucine and tryptophan (–Leu –Trp) for three days. Protein interactions were further examined based on growth on SD medium lacking leucine, tryptophan, histidine (–Leu –Trp –His), as well as lacking leucine, tryptophan, histidine, and adenine (–Leu –Trp –His –Ade). To prevent the leaky expression of *HIS3*, 3-Amino-1,2,4-triazole (3-AT, [BBI, A601149]) was added to the SD medium as suggested by the manufacturer. The detailed procedure has been described in a previous study (Chen et al., 2019).

For dropout assays, 10 μL of the original culture (OD_600_ = 1.0) and diluted cultures (OD_600_ = 0.1, 0.01) were streaked on SD/–Leu –Trp, SD/–Leu –Trp –His supplemented with 3-AT, and SD/–Leu –Trp –His –Ade agar plates as indicated. In the Y2H and Y3H assays, positive transformants were grown on the SD/–Trp –Leu and SD/–Trp –Leu –His for three days, and on the SD/–Trp –Leu –His –Ade for five days at 30°C. Images were acquired with a gel documentation and analysis system (Clnix, GenoSens 2100) using the same parameters. Validation of interactions was conducted through a minimum of three independent assays, and representative images were presented.

### Co-immunoprecipitation and Split-luciferase Complementation Assays

*Agrobacterium tumefaciens* strain GV3101 and *Nicotiana benthamiana* plants were used for the Co-IP and split-LUC assays. The overnight cultures were centrifuged and the pellets were resuspended with the injection buffer (4.43 g/L Murashige and Skoog Basal Medium, 10 mM MES, 10 mM MgCl_2_ and 150 μM Acetosyringone, pH = 5.7). Equal volumes of two adjusted cultures (OD_600_ = 1) were mixed and kept in the dark for 3 h before agroinfiltration. The injected plants were kept in dark overnight and then transferred to normal light conditions.

To conduct the Co-IP assay, total proteins were extracted two days after injection using lysis buffer (50 mM Tris-HCl, pH = 8.0, 150 mM NaCl, 10% glycerol, 1 mM DTT, 1× Cocktail [Yeasen, 20124ES03] and 0.5% NP-40). After centrifugation, one part of the sample was used as input, while the other part was utilized for subsequent experiments. An equal volume of the supernatant was collected and incubated with 15 μL Anti-GFP Nanobody Magarose Beads (AlpalifeBio, KTSM1334) at 4°C for 3 h. The beads were washed with lysis buffer four times and each washing lasted 8 min. Subsequently, the beads were boiled with 2× SDS loading buffer at 95°C for 5 min. The protein samples were separated on 10% SDS-PAGE, and probed with an anti-GFP antibody (1:3000 dilution; Abcam, Ab290) or anti-HA antibody (1:5000 dilution; Yeasen, 30701ES60), followed by the incubation with the correspond secondary antibodies (see “Protein Extraction and Western Blot Analysis” for details).

To perform the split-LUC assay, the injected plants were observed for protein interactions two days after agroinfiltration. The plants were further injected with the solution of 1 mM D-Luciferin potassium salt (Beyotime, ST196) and then incubated for 30 min before observation. The leaves were imaged using a Chemiluminescent Imaging System (Tanon 4600).

### Protein Extraction and Western Blot Analysis

Total protein was extracted from Y2HGold yeast cells according to previous studies (Kushnirov, 2000; Chen et al., 2019). Briefly, three milliliters of yeast cells (OD_600_ = 4.0) were harvested by centrifugation of 13,000 *g*, 1 min at room temperature. The cells were resuspended and pre-treated with 100 μL 2 M LiAc for 5 min on ice. After centrifugation at 1,000 *g* for 1 min, the supernatant was removed and cells were treated with 100 μL 0.4 M NaOH for 5 min on ice. Then, cells were pelleted down by centrifugation at 300 *g* for 2 min and resuspended in 100 μL SDS-PAGE loading buffer (60 mM Tris-HCl pH = 6.8, 10% glycerol, 2% [w/v] SDS, 5% [v/v] β-Mercaptoethanol and 0.0025% [w/v] bromophenol blue). The samples were further boiled at 95°C for 5 min. After centrifugation at 16,000 *g* for 5 min, 20 μL of the supernatant was loaded for each sample.

To extract total protein from rosette leaves of 4- to 5-week-old plants, 50 mg of the leaves were frozen in liquid nitrogen and then homogenized using a high-throughput tissue grinder (Wonbio-48R). The resulting samples were resuspended in 100 μL lysis buffer (50 mM Tris-HCl, 150 mM NaCl, 1 mM EDTA, pH = 8.0, 10% glycerol, 0.2% Triton X-100, and 1× Cocktail) and centrifugated at 16,000 *g* for 5 min at 4°C. The supernatant was boiled at 95°C for 10 min in SDS-PAGE loading buffer (60 mM Tris-HCl pH = 6.8, 10% glycerol, 2% [w/v] SDS, 5% [v/v] β-Mercaptoethanol and 0.0025% [w/v] bromophenol blue). After centrifugation at 16,000 *g* for 5 min, 10 μL of the supernatant was loaded for each sample.

All protein samples were separated by electrophoresis on 10% SDS-PAGE gels and subsequently transferred onto the nitrocellulose membranes (Cytiva, 10600001). After blocking in TBS buffer supplemented with 0.5% (v/v) Tween 20 and 5% (v/v) nonfat milk, the membranes were probed with anti-HA antibody (1:5000 dilution; Yeasen, 30701ES60) or anti-mNeonGreen antibody (1:3000 dilution; Agrisera, AS214525) overnight at 4°C. After washing, the membranes were further incubated with a goat anti-mouse secondary antibody (1:5000 dilution) (Yeasen, 33201ES60), or a goat anti-rabbit secondary antibody (1:5000 dilution) (ABclonal, AS014) for 1 h at room temperature. The protein bands were detected using Super Signal West Pico PLUS Chemiluminescent Substrate (Thermo Fisher Scientific, 34577). Membranes were imaged using a Chemiluminescent Imaging System (Tanon 4600). Regarding Western blot with actin, the primary and secondary antibodies were removed using a mild striping buffer (15 g/L glycine, 1 g/L SDS, 1% tween-20, pH = 2.2), followed by probing the membrane with anti-actin antibody (Abclonal, AC009). Ponceau S (Diamond, A100860) stained nitrocellulose membranes were served as an additional loading control.

### Expression and Observation of Chloroplast Division Proteins in *S. pombe*

Fission yeast *S. pombe* (*h^-^ ade6-210 leu1-32 ura4-D18*) was used in this study. The wild-type yeast cells were cultured in YES medium (5 g/L yeast extract, 30 g/L glucose, and supplemented with 225 mg/L adenine, histidine, leucine, uracil and lysine hydrochloride). Transformation of *S. pombe* cells was conducted using the lithium acetate approach, as previously described (TerBush and Osteryoung, 2012; Chen et al., 2019). Transformed yeast cells were grown on selective synthetic dropout PMG medium (3 g/L potassium hydrogen phthalate, 2.2 g/L Na_2_HPO_4_, 3.75 g/L L-glutamic acid, 20 g/L glucose, 20 mL/L salts, 1 mL/L vitamins, and 0.1 mL/L minerals) lacking either Leu (for pREP41X) or uracil (for pREP42X) or both (for cotransformation) (see https://dornsife.usc.edu/pombenet/media/ for details). Positive colonies expressing the fusion proteins were further verified under microscope, and then inoculated into the corresponding PMG selective media. The cultures were diluted (OD_600_ = 1), and 15 μL of each diluted culture was reinoculated into 2 mL selective PMG media, in order to minimize the growth differences among distinct transformed yeast cells. The cultures were further incubated at 32°C for 36 to 40 h prior to observation.

Transformed cells expressing the chloroplast fusion proteins were visualized by epifluorescence microscopy (Olympus BX53) equipped with a CCD camera (Retiga R6). The fluorescence signals of mCerulean, mVenus and mRuby2 fusion proteins were captured using FluoCa filters FS301 (430 to 450 nm excitation/465 to 495 nm emission), FS303 (480 to 500nm excitation/517.5 to 542.5 nm emission), and FS308 (545 to 575 nm excitation/595 to 665 nm emission), respectively. The excitation intensity of the LED light source (FluoCa) for the mCerulean, mVenus and mRuby2 channel was set to 30%, 5%, and 50%, respectively. The exposure time was 500 ms for all channels. To quantitatively determine the inhibitory effect of ARC3 on FtsZ assembly, the length of the FtsZ1-mVenus filaments (the average length of the two shortest filaments within cells) was measured and normalized to the cell length; the number of the FtsZ1-mVenus filaments was counted manually using Fiji (ImageJ) software (http://fiji.sc/Fiji). The protein expression level of ARC6-mRuby2, ARC6_ΔJLD_-mRuby2 and mRuby2 was assessed by measuring the fluorescence intensity of the mRuby2 signal within cells, which was further normalized to cell size.

### Confocal Microscopy

Localization of ARC3-mNG and ARC6 (or its derivates)-mNG-3×HA fusion proteins in transgenic plants was detected using a laser scanning confocal microscope (Nikon A1 HD25) with a 100× oil immersion objective (numerical aperture, 1.4). Identical confocal setting was applied when observing same fusion protein in distinct transgenic plants.

To detect ARC3-mNG, the abaxial side of leaves from 10- to 14-day-old seedlings was used. The laser intensity for detecting the mNG fluorescence signal (488 nm excitation/500 to 550 nm emission) and the chlorophyll autofluorescence (561 nm excitation/662 to 737 nm emission) was both 10%.

To dissect the localization of ARC6 and its derivates tagged with mNG, leaves from 3- to 4-week-old plants were adopted. The laser intensity for detecting the mNG fluorescence signal (488 nm excitation/500 to 550 nm emission) and the chlorophyll autofluorescence (640 nm excitation/662 to 737 nm emission) was 60 and 55%. Root tips of 7-day-old seedlings and petals of 5-week-old plants were also examined using confocal with the identical setting as leaf samples.

Fluorescent images of immunostained FtsZ2-1 in plants were captured with a laser scanning confocal microscope (Nikon A1 HD25) with a 100× oil immersion objective (numerical aperture, 1.4). To detect Alexa Fluor 488 (488 nm excitation/500 to 550 nm emission) and chlorophyll fluorescence (561 nm excitation/662 to 737 nm emission), the laser intensity was set to 30 and 50%. A 0.5 μm interval was employed to collect z stacks images for both channels. NIS-Elements software was used to perform projection of the obtained z stack images based on the max intensity algorithm.

### Accession Numbers

Sequence data from this study can be found in The Arabidopsis Information Resource (TAIR) database (https://www.arabidopsis.org/) under the following names and accession numbers: *ARC6* (AT5G42480), *ARC3* (AT1G75010), *FtsZ1* (AT5G55280), *FtsZ2-1* (AT2G36250) and *ARC1* (AT4G23940). The single mutants used in this study are *arc6-1* (CS286), *arc6-5* (SAIL_693_G04), *arc3-2* (SALK_057144) and *arc1* (CS262).

**Supplemental Figure 1.** Functional relationship between *ARC6* and *ARC3* during chloroplast division.

**Supplemental Figure 2.** Multiple sequence alignments to identify the conserved tripeptide PPQ in the J-like domain of ARC6.

**Supplemental Figure 3.** The J-like domain of ARC6 does not interact with cpHsp70s.

**Supplemental Figure 4.** Design of constructs for the yeast three-hybrid assays.

**Supplemental Figure 5.** ARC6ΔPPQ promotes the interaction between ARC3 and FtsZs more effectively than ARC6_ΔJLD_.

**Supplemental Figure 6.** Effect of full-length ARC3 on assembly of FtsZ1 filaments in the presence of ARC6 and ARC6_ΔJLD_ in *S. pombe*.

**Supplemental Figure 7.** Effect of ARC6 overexpression in wild-type Col-0.

**Supplemental Figure 8.** Disruption of Z-ring formation by ARC6_ΔJLD_ overexpression depends on ARC3.

**Supplemental Figure 9.** Localization of ARC6-mNG-3×HA in petals and roots.

**Supplemental Figure 10.** Chloroplast and Z-ring morphologies of *arc1* and *arc1 arc6* mutants.

**Supplemental Table 1**. List of primers and restriction enzymes.

## ACKNOWLEDGEMENTS

We thank Xiangwei He and Wenzhu Li from Zhejiang University for providing the *S. pombe* strain and Lu Liu from Shanghai Jiao Tong University for providing the pGREEN vector. We thank the core facilities at the School of Agricultural and Biology, and the School of Life Sciences and Biotechnology from Shanghai Jiao Tong University. This work was supported by the National Science Foundation of China (grant no. 32170333 to C.C.), the Shanghai Pujiang Program from Science and Technology Commission of Shanghai Municipality (grant no. 20PJ1405700 to C.C.), the China Postdoctoral Science Foundation (285751 to L.C. and 278185 to Y.Z.), the Shanghai Super Postdoc Incentive Plan from Shanghai Municipal Human Resources and Social Security Bureau (to L.C.), the U.S. Department of Energy, Office of Science, Basic Energy Sciences (grant no. DE–FG02–06ER15808 to K.W.O.), and the National Science Foundation (grant no. 1719376 to K.W.O.).

## AUTHOR CONTRIBUTIONS

C.C. and K.W.O. conceived the project, and C.C. supervised the study. C.C., W.D. and L.C. designed the experiments; W.D., L.C. and Y.Z. performed most of the experiments and analyzed the data; S.J., M.N., Y.Y., J.M.G. and K.J.P. performed some Y2H assays; Q.H. helped with data analysis; J.X. helped with confocal microscopy; K.W.O. and W.L. provided critical suggestions to the study and the preparation of the manuscript. C.C., L.C., W.D. and K.W.O. wrote the manuscript.

---

**Supplemental Figure 1. Functional relationship between *ARC6* and *ARC3* during chloroplast division.**

(**A-D**) The plant growth phenotype of 3-week-old (**A**) wild-type Col-0, (**B**) *arc3-2*, (**C**) *arc6-5*, and (**D**) *arc3-2 arc6-5*. Scale bars, 2 cm.

**Supplemental Figure 2. Multiple sequence alignments to identify the conserved tripeptide PPQ in the J-like domain of ARC6.**

Protein sequences of DnaJ and J-like domain (JLD) proteins from diverse species were obtained from the National Center for Biotechnology Information (NCBI, https://www.ncbi.nlm.nih.gov/protein/), the Phytozome database at the Joint Genome Institute (JGI, https://phytozome.jgi.doe.gov/pz/portal.html), and Fernbase (https://fernbase.org/). Multiple sequence alignment (MSA) was performed using the European Molecular Biology Laboratory (EMBL)-European Bioinformatics Institute (EBI, https://www.ebi.ac.uk/Tools/msa/muscle/) MUSCLE tools, and the result was submitted to Espript 3.0 (Robert and Gouet, 2014) with the sequence similarities depiction parameter set as %Equivalent and the global score set at 0.7 (http://espript.ibcp.fr/ESPript/ESPript/index.php). Only a partial MSA corresponding to amino acids 89-153 of ARC6 from *Arabidopsis thaliana* is shown. Blue boxes indicate amino acids with 70% or greater identity. Red shading highlights 100% conserved amino acids among all aligned sequences. Asterisks indicate the three conserved residues (PPQ) in angiosperms. The accession numbers of the protein sequences in NCBI are: AAA00009.1 (*Escherichia coli*); AIE75001.1 (*Synechocystis* sp. PCC 6714). The accession number of the protein sequences in Fernbase is Sacu_v1.1_s0076.g017490 (*Salvinia cucullata*). The accession numbers of the protein sequences in JGI are: *Volvox carteri*, Vocar.0016s0119.1.p; *Chlamydomonas reinhardtii*, Cre12.g488500.t1.2; *Marchantia polymorpha*, Mapoly0064s0104.1.p; *Sphagnum fallax*, Sphfalx17G074000.1.p; *Physcomitrella patens*, Pp3c22_12970V3.1.p; *Selaginella moellendorffii*, 92361; *Hordeum vulgare*, HORVU6Hr1G012020.11; *Oryza sativa*, LOC_Os02g03000.2; *Sorghum bicolor*, Sobic.004G020100.1.p; *Zea mays*, Zm00001d053923_P005; *Amborella trichopoda*, evm_27.model.AmTr_v1.0_scaffold00182.17; *Solanum lycopersicum*, Solyc04g081070.3.1; *Solanum tuberosum*, Soltu.DM.04G036190.1; *Theobroma cacao*, Thecc.08G038000.1.p; *Malus domestica*, MD15G1016900; *Vitis vinifera*, VIT_202s0033g00060.1; *Medicago truncatula*, Medtr1g023310.1; *Gossypium raimondii*, Gorai.009G196600.1; *Populus trichocarpa*, Potri.005G235000.4.p; *Glycine max*, Glyma.17G255300.2.p; *Brassica rapa*, Brara.F03845.1.p; *Arabidopsis thaliana*, AT5G42480.1.

**Supplemental Figure 3. The J-like domain of ARC6 does not interact with cpHsp70s.**

Y2H assays of ARC6 J-like domain (ARC6_JLD_) with cpHsp70s. Constructs pGADT7-cpHsp70-1, pGADT7-cpHsp70-2, pGBKT7-ARC6_JLD_, pGBKT7-AtDjA24 were transformed into YH109 yeast cells. Dilutions from the same starting culture are indicated at the bottom. Transformed cells were grown on –Leu –Trp and –Leu –Trp – His medium. The interaction of AtDjA24 with cpHsc70-1 was used as a positive control.

**Supplemental Figure 4. Design of constructs for the yeast three-hybrid assays. (A-E)** Fragments of *ADH1* promoter (fragment 1) and *HA-ARC6_68-614_-ADH1* terminator (fragment 2) were PCR amplified from **(B)** pGADT7-ARC6_68-614_, and subsequently cloned into AvrII-digested pGBKT7 vectors using Gibson Assembly. This process yielded **(C)** pGBKT7-Empty; ARC6_68-614_, **(D)** pGBKT7-FtsZ1; ARC6_68-614_, and **(E)** pGBKT7-FtsZ2; ARC6_68-614_.

**Supplemental Figure 5. ARC6_ΔPPQ_ promotes the interaction between ARC3 and FtsZs more effectively than ARC6_ΔJLD_.**

Y3H assays of the interaction between full-length ARC3 (ARC3_41-741_) and FtsZ proteins in the presence of ARC6_ΔJLD_ or ARC6_ΔPPQ_. Representative images show transformed Y2HGold cells grown on –Leu –Trp and –Leu –Trp –His (200 μM 3-AT) medium for three (left) and four (right) days. Dilutions from the same starting culture are indicated at the bottom.

**Supplemental Figure 6. Effect of full-length ARC3 on assembly of FtsZ1 filaments in the presence of ARC6 and ARC6_ΔJLD_ in *S. pombe*.**

Epifluorescence micrographs of transformed *S. pombe* cells expressing the indicated proteins are shown. Effect of ARC3_41-741_-mCerulean on assembly of FtsZ1-mVenus filaments in strains coexpressing **(A, B)** ARC6_68-614_-mRuby2 and **(C-E)** ARC6_ΔJLD_-mRuby2. **(F)** Effect of ARC3_41-598_-mCerulean on assembly of FtsZ1-mVenus filaments in strains coexpressing mRuby2, which serves as a positive control. In the epifluorescence images, the signals for mCerulean, mVenus, and mRuby2 are falsely colored magenta, green, and yellow, respectively. Scale bars, 5 μm.

**Supplemental Figure 7. Effect of ARC6 overexpression in wild-type Col-0.**

**(A)** Chloroplast morphology (left panels) and FtsZ localization (middle and right panels) in mesophyll cells of T_1_ transgenic plants expressing *35S_pro_:ARC6-mNG-3×HA* in wild-type Col-0. Untransformed wild-type Col-0 is shown in Figure 2A. Chloroplast morphology and FtsZ localization were visualized using DIC microscopy and immunofluorescence staining of FtsZ2-1 (FtsZ), respectively. Scale bars are 20 μm (left panel) and 10 μm (middle and right panels), respectively.

**(B)** Analysis of chloroplast number in relation to mesophyll cell plane area was statistically performed for different genotypes (*n* = 50 cells). The determined slopes for the best-fit lines are: Col-0, 0.018 (R^2^ = 0.95); Col-0 + *35S_pro_:ARC6-mNG-3×HA*, 0.003 (R^2^ = 0.17).

**(C, D)** ARC6 localization in mesophyll cells of Col-0 + *35S_pro_:ARC6-mNG-3×HA* transgenic plants. The fluorescence signals from mNG (green) and chlorophyll autofluorescence (magenta) were captured using confocal laser scanning microscopy. White arrowheads indicated ring-like structures formed by ARC6-mNG-3×HA. Bars are 10 μm.

**(E)** Immunoblot analysis of ARC6-mNG-3×HA fusion protein from T_1_ transgenic Col-0 (lane 4) and *arc3-2* (lane 3) plants expressing *ARC6_pro_:ARC6-mNG-3×HA*. Total proteins extracted from leaf tissue of 4-week-old plants were separated on a 10% SDS-PAGE gel, and the membrane was probed with an anti-HA antibody. The expression of Actin, detected with an anti-Actin antibody, was used as a loading control.

**Supplemental Figure 8. Disruption of Z-ring formation by ARC6_ΔJLD_ overexpression depends on ARC3.**

**(A-C)** Examination of chloroplast morphology (left panels) in 3-week-old mesophyll cells of T_1_ transgenic plants expressing *35S_pro_:ARC6_ΔJLD_-mNG-3×HA* in Col-0 and *arc3-2*. Chloroplast morphology was observed using DIC microscopy. The middle and right panels display merged images of immunostained FtsZ2-1 (green) and chlorophyll autofluorescence (magenta). The inset in the right panel of **(A)** is a magnified image of the mini-ring, as indicated by the dashed line. Immunofluorescence staining images were captured using confocal laser scanning microscopy. Scale bars are as indicated.

**(D)** Immunoblot analysis of the ARC6_ΔJLD_-mNG-3×HA fusion protein in Col-0 + *35S_pro_:ARC6_ΔJLD_-mNG-3×HA* transgenic plants (#2) and *arc3-2* + *35S_pro_:ARC6_ΔJLD_-mNG-3×HA* transgenic plants (#41) is also displayed in Figure 6. The membrane was probed with an anti-HA antibody. Ponceau S-stained large subunit of Rubisco served as a loading control.

**Supplemental Figure 9. Localization of ARC6-mNG-3×HA in petals and roots. (A, B)** ARC6-mNG-3×HA localization in petals from 5-week-old T_1_ *arc6-5* + *ARC6_pro_:ARC6-mNG-3×HA* transgenic plants. **(C, D)** ARC6-mNG-3×HA localization in roots from 7-day-old T_1_ *arc6-5* + *ARC6_pro_:ARC6-mNG-3×HA*. Scale bars are 10 μm for petals and 20 μm for roots. mNG, mNeonGreen; BF, bright field.

**Supplemental Figure 10. Chloroplast and Z-ring morphologies of *arc1* and *arc1 arc6* mutants.**

**(A-D)** Chloroplast morphology (left and middle panels) in 3-week-old mesophyll cells of wild-type L*er*, *arc1*, *arc6-1*, and *arc1 arc6-1* mutant plants. Chloroplast morphology was observed using DIC microscopy. The right panels display merged images of immunostained FtsZ2-1 (green) and chlorophyll autofluorescence (magenta). Scale bars are as indicated.

## REFERENCE

Ajjawi, I., Coku, A., Froehlich, J.E., Yang, Y., Osteryoung, K.W., Benning, C., and Last, R.L. (2011). A J-like protein influences fatty acid composition of chloroplast lipids in Arabidopsis. PLoS One. 6, e25368.

Bruce, B.D. (2000). Chloroplast transit peptides: structure, function and evolution. Trends Cell Biol. 10, 440–447.

Cackett, L., Luginbuehl, L.H., Schreier, T.B., Lopez-Juez, E., and Hibberd, J.M. (2022). Chloroplast development in green plant tissues: the interplay between light, hormone, and transcriptional regulation. New Phytol. 233, 2000–2016.

Chen, C., MacCready, J.S., Ducat, D.C., and Osteryoung, K.W. (2018a). The Molecular Machinery of Chloroplast Division. Plant Physiol. 176, 138–151.

Chen, C., Cao, L., Yang, Y., Porter, K.J., and Osteryoung, K.W. (2019). ARC3 Activation by PARC6 Promotes FtsZ-Ring Remodeling at the Chloroplast Division Site. Plant Cell. 31, 862–885.

Chen, L., Sun, B., Gao, W., Zhang, Q.Y., Yuan, H., and Zhang, M. (2018b). MCD1 Associates with FtsZ Filaments via the Membrane-Tethering Protein ARC6 to Guide Chloroplast Division. Plant Cell. 30, 1807–1823.

Clough, S.J., and Bent, A.F. (1998). Floral dip: a simplified method for Agrobacterium-mediated transformation of Arabidopsis thaliana. Plant J. 16, 735–743.

Colletti, K.S., Tattersall, E.A., Pyke, K.A., Froelich, J.E., Stokes, K.D., and Osteryoung, K.W. (2000). A homologue of the bacterial cell division site-determining factor MinD mediates placement of the chloroplast division apparatus. Curr Biol. 10, 507–516.

Gao, H., Kadirjan-Kalbach, D., Froehlich, J.E., and Osteryoung, K.W. (2003). ARC5, a cytosolic dynamin-like protein from plants, is part of the chloroplast division machinery. Proc.Natl.Acad.Sci.USA 100, 4328–4333.

Gibson, D.G., Young, L., Chuang, R.Y., Venter, J.C., Hutchison, C.A., 3rd, and Smith, H.O. (2009). Enzymatic assembly of DNA molecules up to several hundred kilobases. Nat Methods. 6, 343–345.

Glynn, J.M., Froehlich, J.E., and Osteryoung, K.W. (2008). Arabidopsis ARC6 coordinates the division machineries of the inner and outer chloroplast membranes through interaction with PDV2 in the intermembrane space. Plant Cell. 20, 2460–2470.

Glynn, J.M., Miyagishima, S.Y., Yoder, D.W., Osteryoung, K.W., and Vitha, S. (2007). Chloroplast division. Traffic. 8, 451–461.

Glynn, J.M., Yang, Y., Vitha, S., Schmitz, A.J., Hemmes, M., Miyagishima, S.Y., and Osteryoung, K.W. (2009). PARC6, a novel chloroplast division factor, influences FtsZ assembly and is required for recruitment of PDV1 during chloroplast division in Arabidopsis. Plant J. 59, 700–711.

Johnson, C.B., Shaik, R., Abdallah, R., Vitha, S., and Holzenburg, A. (2015). FtsZ1/FtsZ2 Turnover in Chloroplasts and the Role of ARC3. Microsc Microanal. 21, 313–323.

Johnson, C.B., Tang, L.K., Smith, A.G., Ravichandran, A., Luo, Z., Vitha, S., and Holzenburg, A. (2013). Single particle tracking analysis of the chloroplast division protein FtsZ anchoring to the inner envelope membrane. Microsc Microanal. 19, 507–512.

Kadirjan-Kalbach, D.K., Yoder, D.W., Ruckle, M.E., Larkin, R.M., and Osteryoung, K.W. (2012). FtsHi1/ARC1 is an essential gene in Arabidopsis that links chloroplast biogenesis and division. Plant J. 72, 856–867.

Keeling, P.J. (2010). The endosymbiotic origin, diversification and fate of plastids. Philos Trans R Soc Lond B Biol Sci. 365, 729–748.

Kohler, R.H., Cao, J., Zipfel, W.R., Webb, W.W., and Hanson, M.R. (1997). Exchange of protein molecules through connections between higher plant plastids. Science. 276, 2039–2042.

Kushnirov, V.V. (2000). Rapid and reliable protein extraction from yeast. Yeast. 16, 857–860.

Lee, D.W., and Hwang, I. (2018). Evolution and Design Principles of the Diverse Chloroplast Transit Peptides. Mol Cells. 41, 161–167.

Leech R, Baker NR (1983) The development of photosynthetic capacity in leaves. In JE Dale, FL Milthorpe, eds, The Growth and Functioning of Leaves. Cambridge University Press, Cambridge, pp 271–307

Liu, X., An, J., Wang, L., Sun, Q., An, C., Wu, B., Hong, C., Wang, X., Dong, S., Guo, J., Feng, Y., and Gao, H. (2022). A novel amphiphilic motif at the C-terminus of FtsZ1 facilitates chloroplast division. Plant Cell. 34, 419–432.

Maple, J., Chua, N.H., and Moller, S.G. (2002). The topological specificity factor AtMinE1 is essential for correct plastid division site placement in Arabidopsis. Plant J. 31, 269–277.

Maple, J., Aldridge, C., and Moller, S.G. (2005). Plastid division is mediated by combinatorial assembly of plastid division proteins. Plant J. 43, 811–823.

Maple, J., Vojta, L., Soll, J., and Moller, S.G. (2007). ARC3 is a stromal Z-ring accessory protein essential for plastid division. EMBO Rep. 8, 293–299.

Martin, W.F., Garg, S., and Zimorski, V. (2015). Endosymbiotic theories for eukaryote origin. Philos Trans R Soc Lond B Biol Sci. 370, 20140330.

McAndrew, R.S., Olson, B.J., Kadirjan-Kalbach, D.K., Chi-Ham, C.L., Vitha, S., Froehlich, J.E., and Osteryoung, K.W. (2008). In vivo quantitative relationship between plastid division proteins FtsZ1 and FtsZ2 and identification of ARC6 and ARC3 in a native FtsZ complex. Biochem J. 412, 367–378.

McKinnon, L., and Theg, S.M. (2019). Determinants of the Specificity of Protein Targeting to Chloroplasts or Mitochondria. Mol Plant. 12, 893–895.

McQuillen, R., and Xiao, J. (2020). Insights into the Structure, Function, and Dynamics of the Bacterial Cytokinetic FtsZ-Ring. Annu Rev Biophys. 49, 309–341.

Miyagishima, S., Takahara, M., Mori, T., Kuroiwa, H., Higashiyama, T., and Kuroiwa, T. (2001). Plastid division is driven by a complex mechanism that involves differential transition of the bacterial and eukaryotic division rings. Plant Cell. 13, 2257–2268.

Miyagishima, S.Y., Froehlich, J.E., and Osteryoung, K.W. (2006). PDV1 and PDV2 mediate recruitment of the dynamin-related protein ARC5 to the plastid division site. Plant Cell. 18, 2517–2530.

Miyagishima, S.Y., Nishida, K., Mori, T., Matsuzaki, M., Higashiyama, T., Kuroiwa, H., and Kuroiwa, T. (2003). A plant-specific dynamin-related protein forms a ring at the chloroplast division site. Plant Cell. 15, 655–665.

Nakanishi, H., Suzuki, K., Kabeya, Y., and Miyagishima, S.Y. (2009). Plant-specific protein MCD1 determines the site of chloroplast division in concert with bacteria-derived MinD. Curr Biol. 19, 151–156.

Olson, B.J., Wang, Q., and Osteryoung, K.W. (2010). GTP-dependent heteropolymer formation and bundling of chloroplast FtsZ1 and FtsZ2. J Biol Chem. 285, 20634–20643.

Osteryoung, K.W., and Pyke, K.A. (2014). Division and dynamic morphology of plastids. Annu Rev Plant Biol. 65, 443–472.

Osteryoung, K.W., Stokes, K.D., Rutherford, S.M., Percival, A.L., and Lee, W.Y. (1998). Chloroplast division in higher plants requires members of two functionally divergent gene families with homology to bacterial *ftsZ*. Plant Cell. 10, 1991–2004.

Porter, K.J., Cao, L., Chen, Y., TerBush, A.D., Chen, C., Erickson, H.P., and Osteryoung, K.W. (2021). The *Arabidopsis thaliana* chloroplast division protein FtsZ1 counterbalances FtsZ2 filament stability *in vitro*. J Biol Chem. 296, 100627.

Pulido, P., and Leister, D. (2018). Novel DNAJ-related proteins in *Arabidopsis thaliana*. New Phytol. 217, 480–490.

Pyke, K.A., and Leech, R.M. (1991). Rapid Image Analysis Screening Procedure for Identifying Chloroplast Number Mutants in Mesophyll Cells of *Arabidopsis thaliana* (L.) Heynh. Plant Physiol. 96, 1193–1195.

Pyke, K.A., and Leech, R.M. (1992). Chloroplast Division and Expansion Is Radically Altered by Nuclear Mutations in *Arabidopsis thaliana*. Plant Physiol. 99, 1005–1008.

Pyke, K.A., Rutherford, S.M., Robertson, E.J., and Leech, R.M. (1994). *arc6*, A Fertile Arabidopsis Mutant with Only Two Mesophyll Cell Chloroplasts. Plant Physiol. 106, 1169–1177.

Schmitz, A.J., Glynn, J.M., Olson, B.J., Stokes, K.D., and Osteryoung, K.W. (2009). *Arabidopsis* FtsZ2-1 and FtsZ2-2 are functionally redundant, but FtsZ-based plastid division is not essential for chloroplast partitioning or plant growth and development. Mol Plant. 2, 1211–1222.

Shimada, H., Koizumi, M., Kuroki, K., Mochizuki, M., Fujimoto, H., Ohta, H., Masuda, T., and Takamiya, K. (2004). ARC3, a chloroplast division factor, is a chimera of prokaryotic FtsZ and part of eukaryotic phosphatidylinositol-4-phosphate 5-kinase. Plant Cell Physiol. 45, 960–967.

Sung, M.W., Shaik, R., TerBush, A.D., Osteryoung, K.W., Vitha, S., and Holzenburg, A. (2018). The chloroplast division protein ARC6 acts to inhibit disassembly of GDP-bound FtsZ2. J Biol Chem. 293, 10692–10706.

Tamadaddi, C., Verma, A.K., Zambare, V., Vairagkar, A., Diwan, D., and Sahi, C. (2022). J-like protein family of *Arabidopsis thaliana*: the enigmatic cousins of J-domain proteins. Plant Cell Rep 41, 1343–1355.

TerBush, A.D., and Osteryoung, K.W. (2012). Distinct functions of chloroplast FtsZ1 and FtsZ2 in Z-ring structure and remodeling. J Cell Biol. 199, 623–637.

TerBush, A.D., Yoshida, Y., and Osteryoung, K.W. (2013). FtsZ in chloroplast division: structure, function and evolution. Curr Opin Cell Biol. 25, 461–470.

TerBush, A.D., Porzondek, C.A., and Osteryoung, K.W. (2016). Functional Analysis of the Chloroplast Division Complex Using *Schizosaccharomyces pombe* as a Heterologous Expression System. Microsc Microanal. 22, 275–289.

TerBush, A.D., MacCready, J.S., Chen, C., Ducat, D.C., and Osteryoung, K.W. (2018). Conserved Dynamics of Chloroplast Cytoskeletal FtsZ Proteins Across Photosynthetic Lineages. Plant Physiol. 176, 295–306.

Vitha, S., and Osteryoung, K.W. (2011). Immunofluorescence microscopy for localization of Arabidopsis chloroplast proteins. Methods Mol Biol. 774, 33–58.

Vitha, S., McAndrew, R.S., and Osteryoung, K.W. (2001). FtsZ ring formation at the chloroplast division site in plants. J Cell Biol. 153, 111–120.

Vitha, S., Froehlich, J.E., Koksharova, O., Pyke, K.A., van Erp, H., and Osteryoung, K.W. (2003). ARC6 is a J-domain plastid division protein and an evolutionary descendant of the cyanobacterial cell division protein Ftn2. Plant Cell. 15, 1918–1933.

Wang, W., Li, J., Sun, Q., Yu, X., Zhang, W., Jia, N., An, C., Li, Y., Dong, Y., Han, F., Chang, N., Liu, X., Zhu, Z., Yu, Y., Fan, S., Yang, M., Luo, S.Z., Gao, H., and Feng, Y. (2017). Structural insights into the coordination of plastid division by the ARC6-PDV2 complex. Nat Plants. 3, 17011.

Yoder, D.W., Kadirjan-Kalbach, D., Olson, B.J., Miyagishima, S.Y., Deblasio, S.L., Hangarter, R.P., and Osteryoung, K.W. (2007). Effects of mutations in Arabidopsis FtsZ1 on plastid division, FtsZ ring formation and positioning, and FtsZ filament morphology in vivo. Plant Cell Physiol. 48, 775–791.

Yoshida, Y. (2018). Insights into the Mechanisms of Chloroplast Division. Int J Mol Sci. 19.

Yoshida, Y., Mogi, Y., TerBush, A.D., and Osteryoung, K.W. (2016). Chloroplast FtsZ assembles into a contractible ring via tubulin-like heteropolymerization. Nat Plants. 2, 16095.

Zhang, M., Schmitz, A.J., Kadirjan-Kalbach, D.K., Terbush, A.D., and Osteryoung, K.W. (2013). Chloroplast division protein ARC3 regulates chloroplast FtsZ-ring assembly and positioning in arabidopsis through interaction with FtsZ2. Plant Cell. 25, 1787–1802.

Zhang, M., Chen, C., Froehlich, J.E., TerBush, A.D., and Osteryoung, K.W. (2016). Roles of Arabidopsis PARC6 in Coordination of the Chloroplast Division Complex and Negative Regulation of FtsZ Assembly. Plant Physiol. 170, 250–262.

Zhang, Y., Zhang, X., Cui, H., Ma, X., Hu, G., Wei, J., He, Y., and Hu, Y. (2021). Residue 49 of AtMinD1 Plays a Key Role in the Guidance of Chloroplast Division by Regulating the ARC6-AtMinD1 Interaction. Front Plant Sci. 12, 752790.

Zimorski, V., Ku, C., Martin, W.F., and Gould, S.B. (2014). Endosymbiotic theory for organelle origins. Curr Opin Microbiol. 22, 38–48.

